# A porcine chronic hepatitis E virus (HEV) infection model exhibits HEV replication in male accessory reproductive glands and immune-mediated reproductive damage

**DOI:** 10.1101/2024.07.10.602840

**Authors:** Kush K. Yadav, Patricia A Boley, Thamonpan Laocharoensuk, Saroj Khatiwada, Carolyn M Lee, Menuka Bhandari, Juliette Hanson, Scott P. Kenney

## Abstract

Hepatitis E virus (HEV) is an expanding zoonotic viral disease threat. Although HEV causes acute viral hepatitis, it is increasingly being recognized as a systemic pathogen with detection and damage in extrahepatic tissues. The presence of HEV RNA in the semen of chronically infected human patients in the absence of viremia and fecal shedding and presence of HEV in the sperm head underscores the need to understand the interaction of HEV within the male reproduction system. Male accessory glands secrete biofluids necessary for sperm nourishment and to neutralize the acidity of the vagina. The role of male accessory glands in the dissemination and persistence of HEV infection have not been studied. Using an immunosuppressed pig model for chronic HEV infection, we demonstrate infectious HEV in mature sperm cells altering the sperm motility and morphology. HEV isolated from sperm cells remained infectious in human hepatoma cells. Spermatic fluid contained lower virus titers than the sperm cells from chronically infected pigs highlighting that the sperm cells themselves can associate with the virus. Evaluation of the male accessory glands demonstrated viral replication, infiltration of CD45 leukocytes, and apoptosis associated with HEV infection. A decrease in serum testosterone levels was evident in the HEV infected pigs. Even though a lower viral RNA titer was seen in serum and feces of chronically infected, immunosuppressed and ribavirin treated pigs, high viral RNA and infectious particles in sperm is a concern. Our findings necessitate further studies defining the mechanism of sperm cell invasion by HEV, length of HEV survival in sperm cells during chronic HEV infection, and risk of sexual transmission of HEV during both acute and chronic phases of infection.

**Author Summary:** Hepatitis E virus, a leading cause of acute viral hepatitis, causes both acute and chronic infection in humans. Recent advances within the HEV field have demonstrated extrahepatic diseases associated with HEV. More recent findings have revealed infectious HEV in the vagina, Sertoli cells, and ejaculate of humans, and sperm cells of pigs. We demonstrate that the male accessory sex glands may have a role in the persistence of HEV infection during chronic infections. We utilized an established immunosuppressed pig model and treated pigs with ribavirin to study the presence of virus in the sperm cells. We demonstrated high viral RNA loads and infectious particles associated with sperm cells. Our study further highlights the importance of the testis, as an immune privileged site, in the maintenance of chronic HEV infection. New studies to evaluate the mechanisms by which HEV associates with sperm cells, the length of HEV survival in sperm cell fractions, and consideration of the testes as a potential HEV reservoir are necessary.

## Introduction

Hepatitis E virus (HEV) is a major challenge for medical and scientific communities. Among the many issues complicating HEV disease are increasingly complicated extrahepatic manifestations (1). These extrahepatic manifestations are most frequently encountered with immunosuppressed patients where the HEV infection progresses to a chronic state (2–4). HEV in tissues beyond the liver and its association with the disease have shifted the study of hepevirus-host interactions, expanding the need to examine sites outside of the liver to fully understand HEV pathogenesis.

Recently HEV RNA has been detected in the semen of seven chronically infected human patients (5) and in human semen samples where the presence of HEV RNA correlated with infertility (6). Interestingly, in one of the chronically infected patients, viremia and fecal virus shedding had subsided with only semen demonstrating HEV RNA (5). The seminal HEV RNA concentration was reported to be 100-fold higher compared to the serum among the five other chronically infected patients (7). Previously, we defined HEV association with the sperm head altering sperm quality in pigs (8). We found infectious virus associated with sperm cells when tested in vitro. We demonstrated higher HEV titers from sperm cell fractions than in the seminal fluid using a natural HEV host, pigs (8). Our initial findings demonstrated that the pig could be a good reproductive model to assess HEV effects in male reproductive disorders (8).

Human male reproductive accessory glands such as the prostate gland, seminal vesicle, and Cowper’s gland are also found in pigs allowing the pig to be used as a model to study the male accessory glands (9). Male accessory gland infection (MAGI) is known to compromise male fertility via production of reactive oxygen species (ROS), impaired secretory capacity of the accessory glands, and via the direct effect of microorganisms on spermatozoa (10, 11). Literature on the most studied sexually transmittable disease (STD), human immunodeficiency virus (HIV), suggests the prostate gland (12) is a primary tissue reservoir helping in the persistence of the virus and resulting in transmission via coitus. The inflammatory response to viral infection in the male accessory glands has been demonstrated to alter sperm viability parameters (11, 13). Prostate gland inflammation is also directly associated with altered concentrations of testosterone (14). Other hepatitis causing viruses such as hepatitis B virus (HBV) and hepatitis C virus (HCV) have been shown to infect the testes and have been correlated to male infertility. HEV effects in the male reproductive system have been correlated to infertility and testicular damage using animal models such as rhesus macaques (6), mice (15), and Mongolian gerbils (16). However, the role of male accessory glands in the contribution of HEV in the sperm cells and seminal fluids have not been studied. Hence, we sought to understand the role of male accessory glands in HEV replication and persistence.

Even though HEV has been described as self-limiting illness, chronic clinical manifestation has been observed in immunosuppressed individuals, particularly organ transplant patients, HIV patients, and cancer patients (17). A majority of the chronic cases are due to genotype (gt) 3 and gt4 *Paslahepevirus balayani* strains (2, 18). Studies show 10% of chronically infected patients have higher chances of developing cirrhosis within 2-5 years (19). The first step in treating chronic HEV infection in solid organ transplant patients is to reduce the immunosuppressive drugs followed by the administration of ribavirin (RBV) (17, 20). It has been documented that the achieved sustained virological response (SVR) rate is 91%, 76%, 67%, and 63% in the liver, kidney, lung, and heart transplant patients, respectively (17). Additionally, no response was documented in 6% of the chronically infected patients and relapse has been shown in 18% of the chronically infected patients (17, 19). The detection of HEV in the ejaculate of seven out of nine chronically infected men highlights the need to understand the reproductive tissue’s role in the maintenance of HEV infection (7, 21).

The lower SVR rate and relapse in the chronically infected patients raises the concern about the potential immune privileged tissue reservoir, such as testis role in the persistence or relapse of chronic HEV infection. To mimic the immunosuppressed chronically infected human patient, we utilized an established chronic HEV pig model (22) to understand the effect of HEV in sperm quality and morphology. We further studied the effect of HEV on blood-testis-barrier (BTB) destruction, infiltration of leukocytes into the testis, testosterone concentration and contribution of male accessory glands in the persistence of HEV during chronic HEV infection. Answers to these questions are important to understand the tissue reservoirs of HEV in the male reproductive system and the ability of the virus to transmit sexually between partners.

## Results

### HEV is associated with the head region of sperm cells

In this study, we utilized the immunosuppressed pig model for chronic HEV infection (23). Immunosuppressed pigs chronically infected with the *Paslahepevirus balayani* genotype 3 US-2 strain of HEV demonstrated viremia, fecal virus shedding with low HEV titers until the study was terminated on day 84 (Fig. 1A). Sperm cells separated from the seminal fluid of the US-2 HEV infected immunosuppressed pigs demonstrated higher viral titers than that present in seminal fluid, serum, or feces (Fig. 1B). Using flow cytometry analysis, we report that at least 7% of the sperm cells from the chronically infected, immunosuppressed and ribavirin treated pigs contained HEV antigen (Fig. 1C). Sperm cells collected from the immunosuppressed and ribavirin treated pigs demonstrated the presence of HEV ORF2 antigen associated with the sperm head (acrosomal region) while the uninfected mock group demonstrated no HEV staining (Fig. 1D).

**Figure 1.**
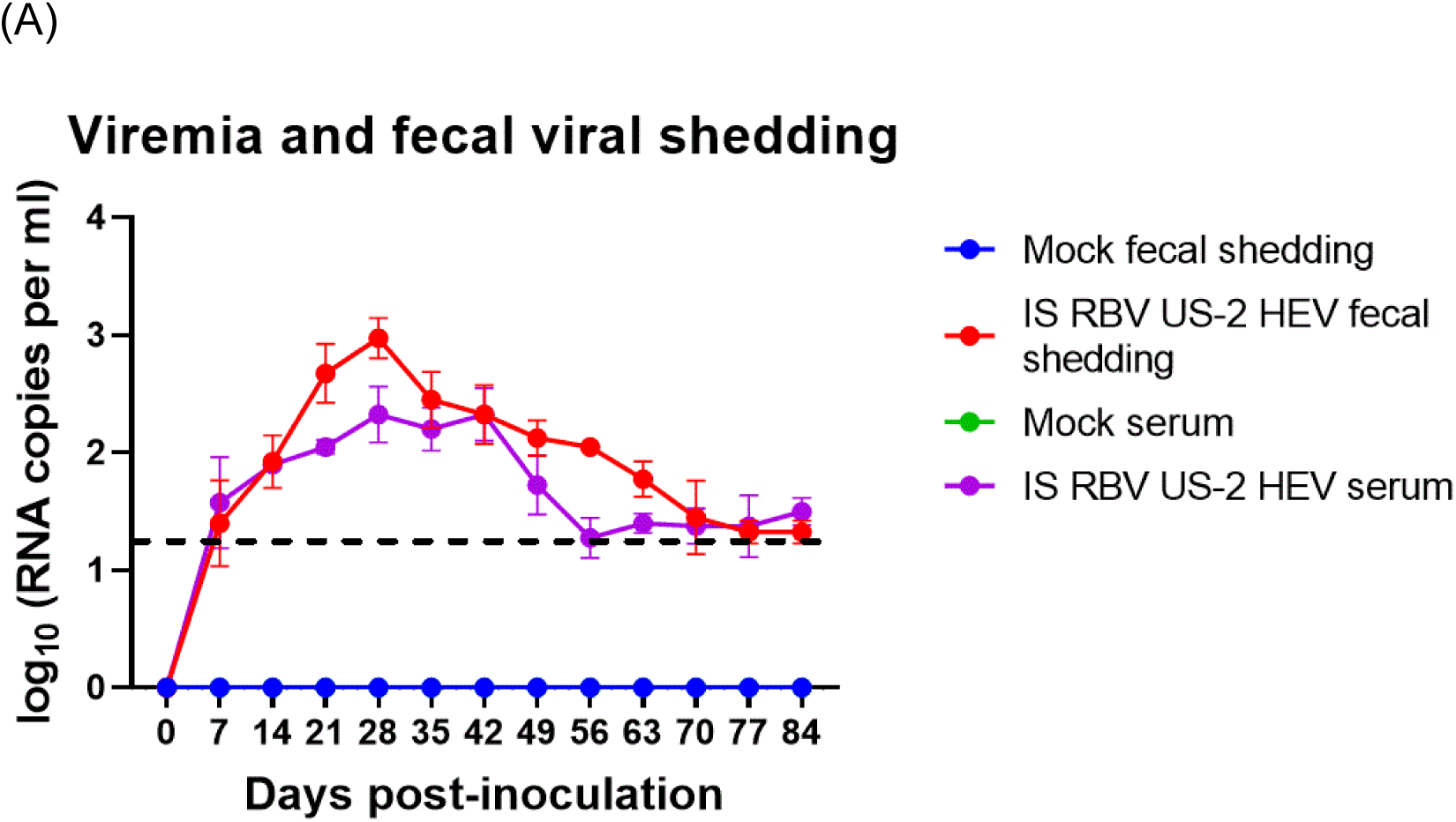
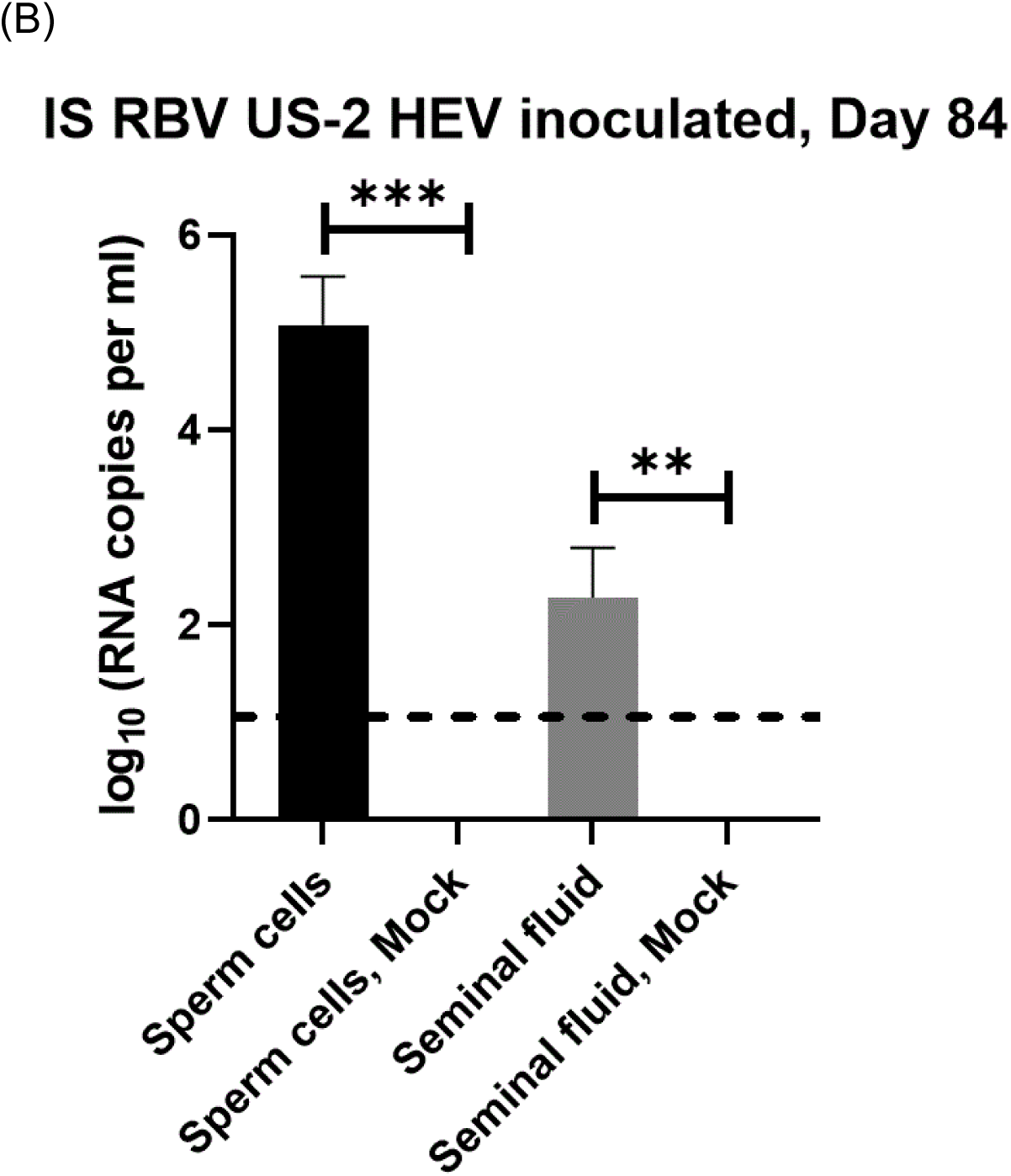
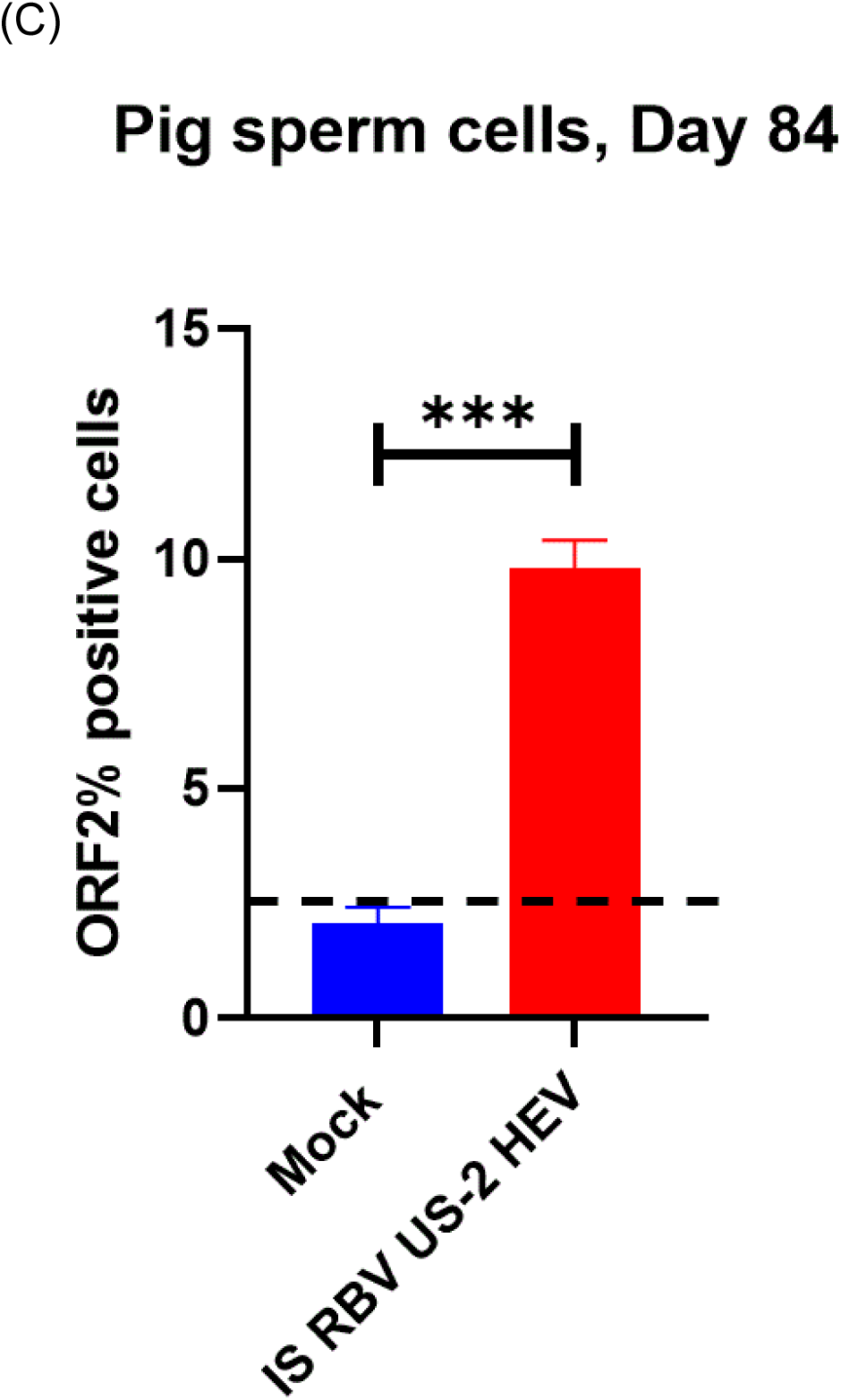
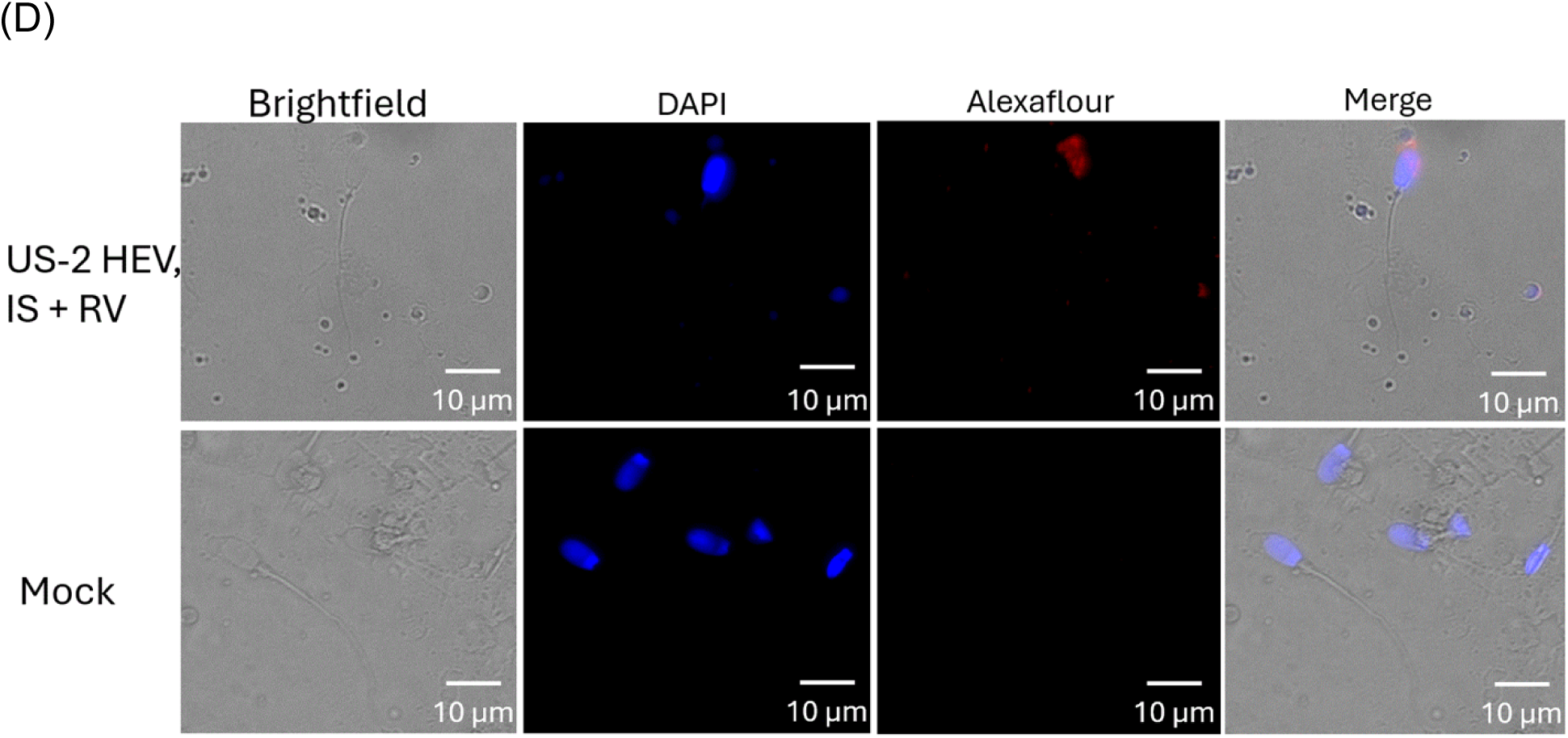
US-2 HEV infected, immunosuppressed (IS) and ribavirin (RBV) treated sperm cells associate with hepatitis E virus. (A) Viremia and fecal viral shedding from IS, RBV treated pigs inoculated with US-2 HEV. Mock-infected groups remained negative throughout the study. (B) HEV RNA loads in sperm cells suspension and spermatic fluid from IS RBV treated pigs inoculated with US-2 HEV. (C) Flow cytometry analysis showing the percentage of sperm cells containing hepatitis E virus (US-2 strain). (D) Immunohistochemical detection of hepatitis E (red) in the acrosomal region of sperm head obtained from the IS RBV US-2 HEV infected pigs at day 84 post infection; HEV open reading frame (ORF2) is red; DAPI stain is blue (nucleus). The dotted line represents the cut-off value demonstrating the background. ** indicates p < 0.01, *** indicates p < 0.001, **** indicates p < 0.0001.

### Sperm cells collected from immunosuppressed US-2 HEV infected pigs are infectious to Huh7 S10-3 cells

Higher HEV RNA copies were seen in sperm cell lysates in comparison to the seminal fluid (Fig.1B). To evaluate the infectious ability of the HEV associated with the sperm cells, we lysed sperm (∼6.2 × 10^5^ viral RNA copies) cells and inoculated human liver cells (Huh7 S10-3) *in vitro*. HEV ORF2 in Huh7 S10-3 cells was detected 5 days post inoculation with sperm cell lysates demonstrating the HEV associated with the sperm cells was viable virus (Fig. 2A). Inoculated HEV was further shown to be replication competent by an increase in viral RNA in supernatant harvested from the inoculated Huh7 S10-3 cells from day 0, 3, and 5 (Fig. 2B).

**Figure 2.**
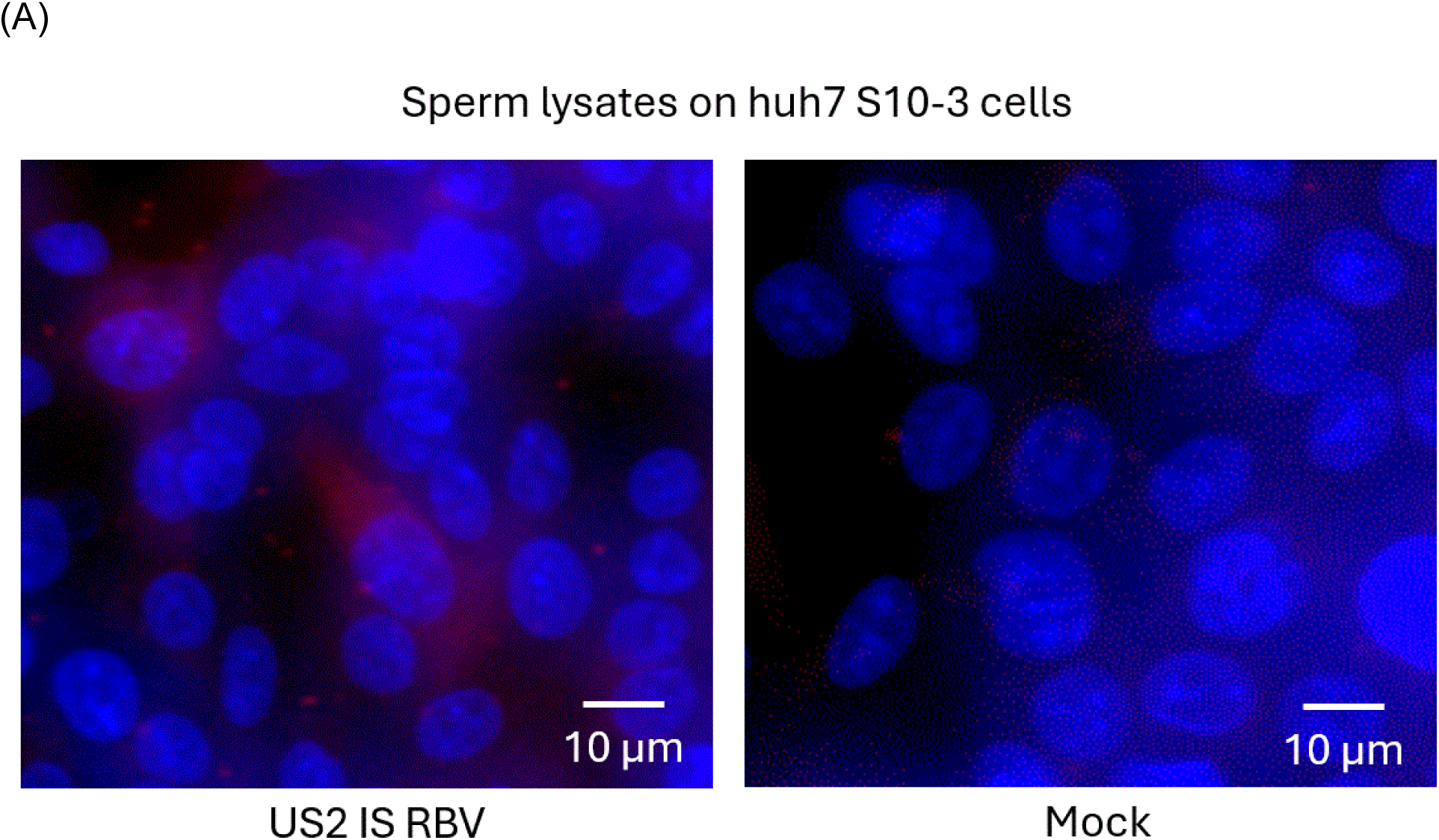
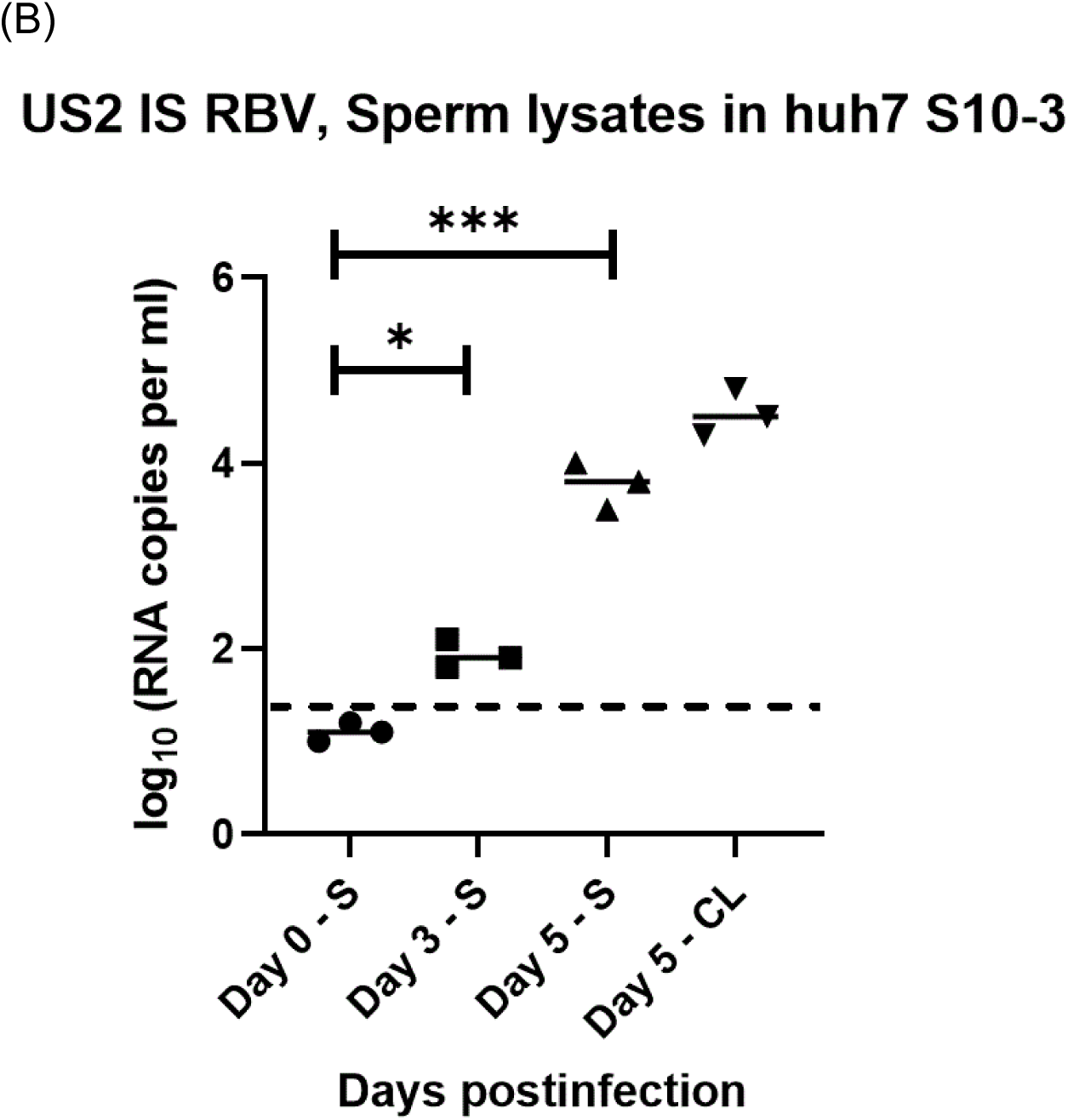
Sperm cells collected from the US-2 HEV infected, immunosuppressed (IS) and ribavirin (RBV) treated pigs are infectious to Huh7 s10-3 cells. (A) Immunodetection of hepatitis E showing infectious sperm in the Huh7 S10-3 cells. (B) Replication kinetics of sperm derived HEV using Huh7 S10-3 cells. US-2 HEV RNA loads in culture supernatant (S) and cell lysates (CL) of Huh7 S10-3 cell cultures after inoculation with the lysed sperm cells collected from the IS RBV US-2 HEV inoculated pigs. Independent biological experiments, mean ± SD of three replicates, are presented. The dotted line represents the cut-off value demonstrating the background from initial attachment of the virus to the cell surfaces. ** indicates p < 0.01, *** indicates p < 0.001.

### Sperm motility and morphology were altered by HEV infection in the immunosuppressed pigs

Semen analysis was immediately performed after collection of semen from the epididymis. To characterize the motility and morphological difference between groups, 200 sperm cells were analyzed. The progressive sperm motility in the immunosuppressed US-2 HEV infected pigs (PR% = 63% ± 2%) was found to be significantly lower than the mock infected pig sperm (78% ± 3%) and immunosuppressed, ribavirin treated (77% ± 2%) pigs (Fig. 3A). The non-progressive sperm motility in US-2 HEV infected pigs (NP% = 20% ± 5%) was found to be significantly higher than the mock infected (12% ± 2%) and immunosuppressed, ribavirin treated (10% ± 4%) pigs (Fig. 3A). Immobility of sperm was higher in sperm from the US-2 HEV infected pigs (21% ± 7%) when compared to mock infected (10% ± 3%) and immunosuppressed, ribavirin treated (13% ± 4%) pigs (Fig. 3A). Morphologically, we found a higher percentage of abnormal sperm heads (40% ± 4%) and tails (15% ± 3%) in US-2 HEV infected pigs (Fig. 3B).

**Figure 3.**
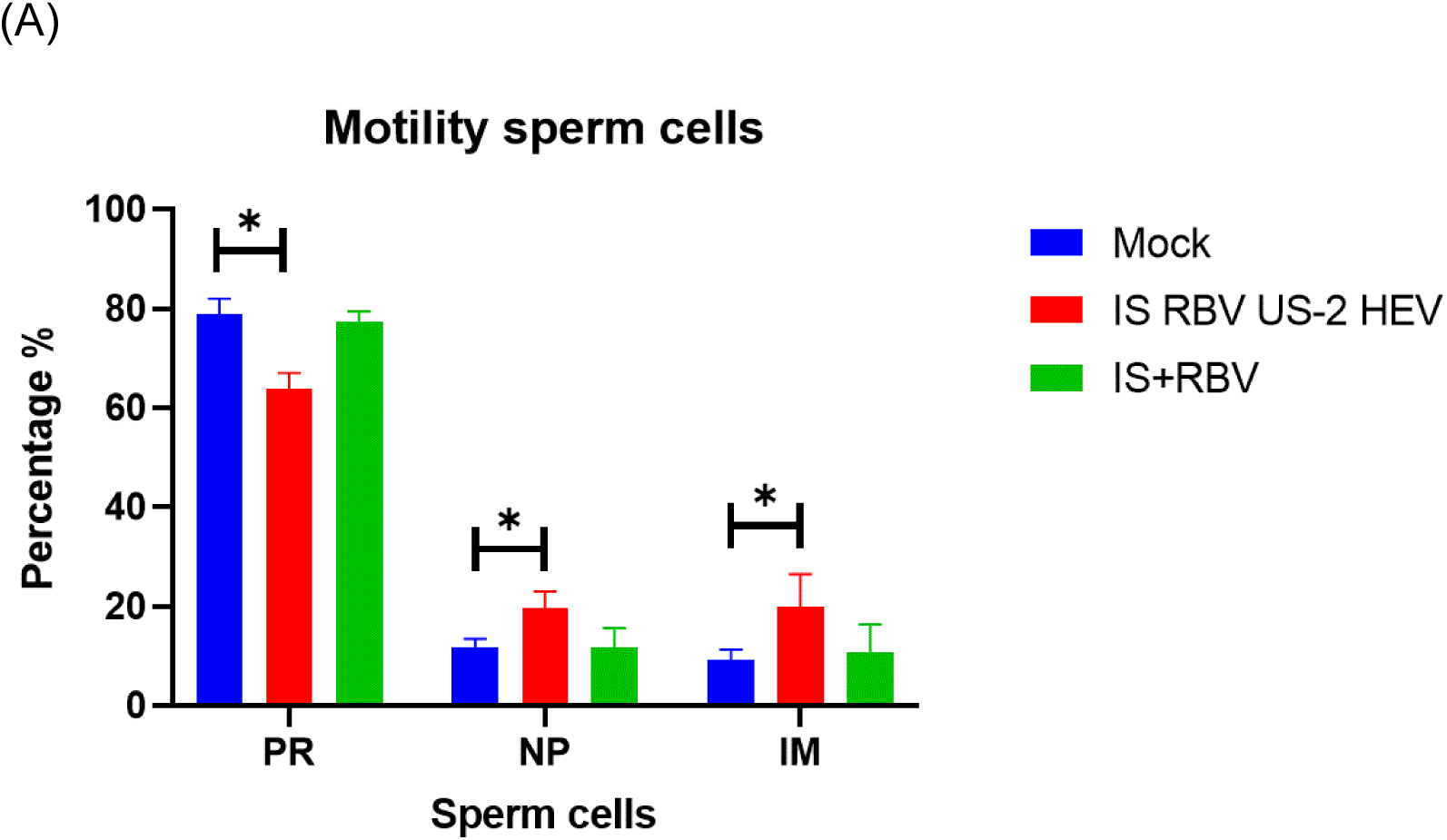
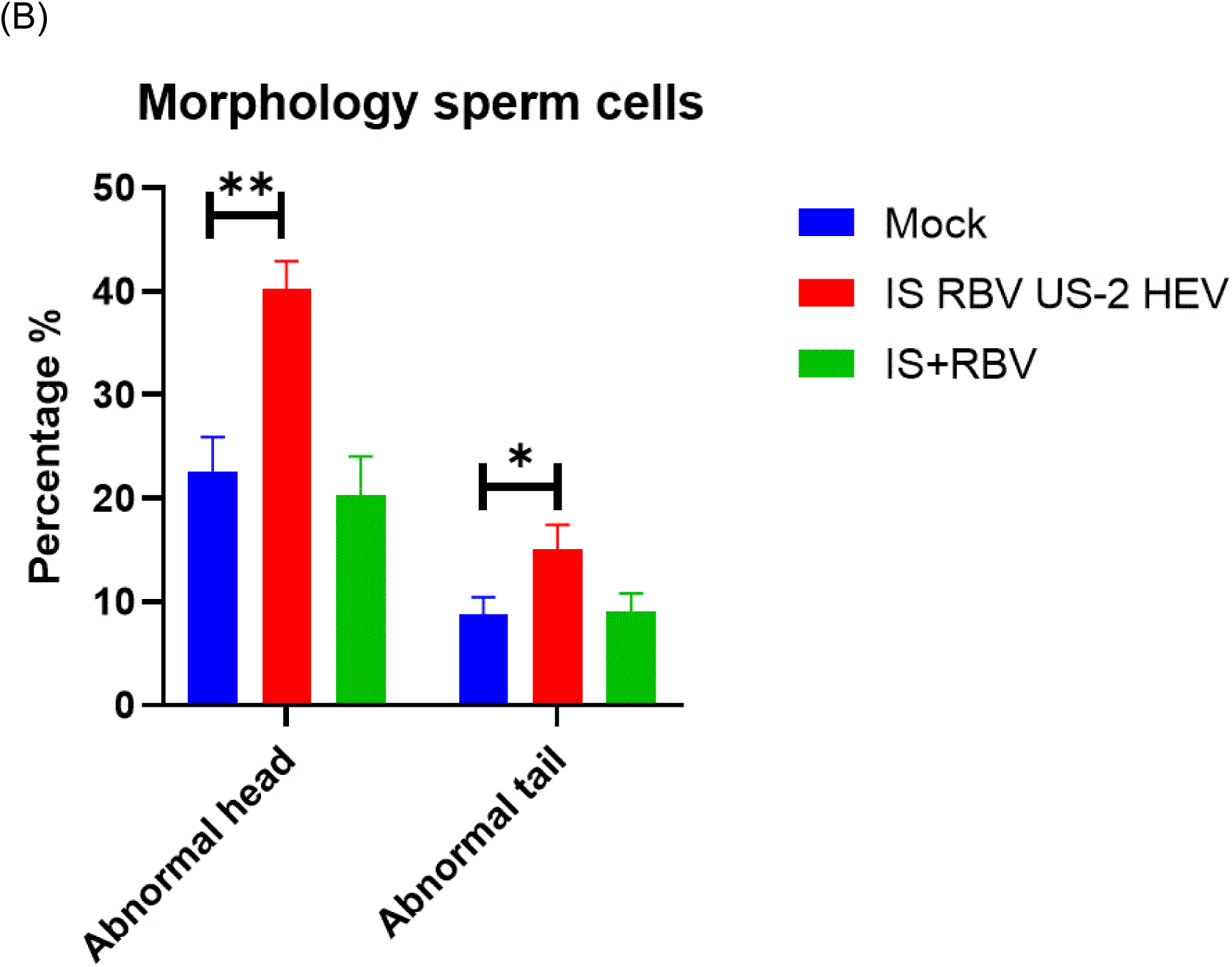
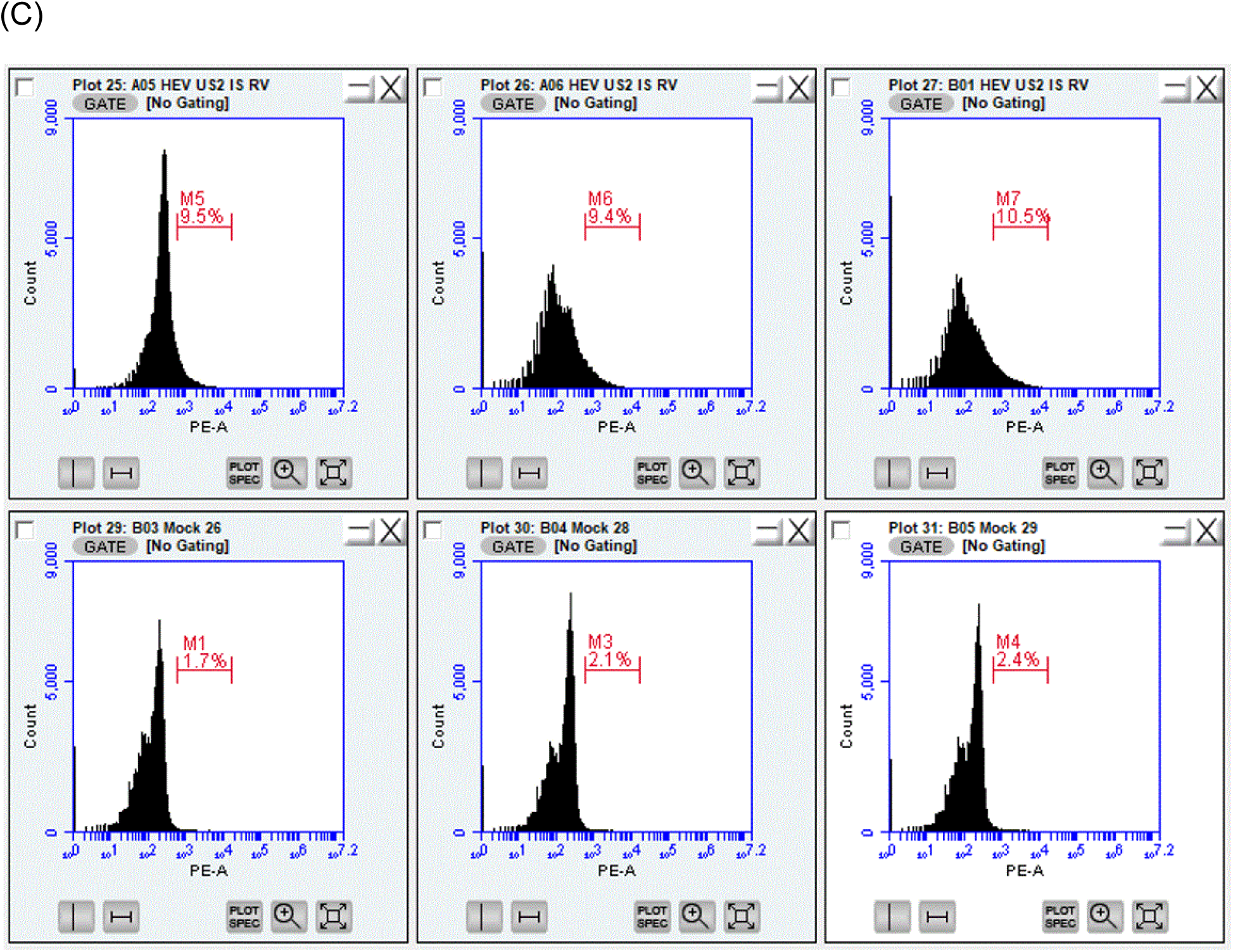
Hepatitis E virus alters mature sperm cell motility and morphology in immunosuppressed (IS), ribavirin (RBV) treated pigs. (A) Light microscopic observation of 200 live mature sperm cells harvested from mock, IS + RBV, or IS, RBV and infected pig epididymis. Sperm cells demonstrated decreased progressive motility when infected by HEV US-2 in IS, RBV treated group. PR – progressive motility of sperm (moving active, either linearly or in a circle, regardless of speed); NP – non-progressive motility (all other patterns of motility with absent progression). IM – immobility. (B) Light microscopic observation of live sperm cells harvested from mock, IS + RBV, or IS, RBV and US-2 HEV infected pig epididymis. Sperm from US-2 HEV infected IS, RBV treated pigs showed a significant increase in mature sperm cells with head abnormalities. No significant changes were observed in the tail of the sperm cells. * indicates p < 0.05, ** indicates p < 0.01. (C) A histogram plot was used to show the flow cytometry results. Sperm cells from mock non-infected pigs and from US-2 HEV infected, immunosuppressed (IS) and ribavirin (RBV) treated pigs.

### HEV infection leads to testicular damage through apoptosis and induces a hormonal disorder

The blood testis barrier (BTB) protects the testis against invading pathogens and restricts immune cell entry to prevent immune-mediated sperm cell damage (24) Apoptosis of the Sertoli cells forming the BTB leads to inflammatory changes and eventually causes a disturbance in the spermatogonia (25). Immunohistochemistry of the testicles was performed to explore the mechanisms underlying the reductions in the sperm quality caused by HEV infection. HEV ORF2 antigen was observed at the BTB of pigs harvested at 84 days post inoculation (Fig. 4A). The breakage of BTB caused by HEV infection may have accounted for the increased infiltration of the testis of HEV infected pigs by CD45+ leukocytes seen at 84 dpi (Fig. 4B). Terminal deoxynucleotidyl transferase biotin-dUTP nick end labeling (TUNEL) assays revealed the apoptosis of cells bordering the BTB (Fig. 4C). Serum testosterone concentrations were significantly decreased in US-2 HEV infected pigs tested on day 70 and 84 post inoculation compared to mock pigs (Fig. 4D). Interestingly, pigs under immunosuppressive and ribavirin drug treatment also demonstrated a decrease in the serum testosterone concentration similar to the US-2 HEV infected, immunosuppressed and ribavirin treated pigs (Fig. 4E). However, on day 84, a significantly lower level of testosterone was observed only in US-2 HEV infected, immunosuppressed and ribavirin treated pigs (Fig. 4E).

**Figure 4.**
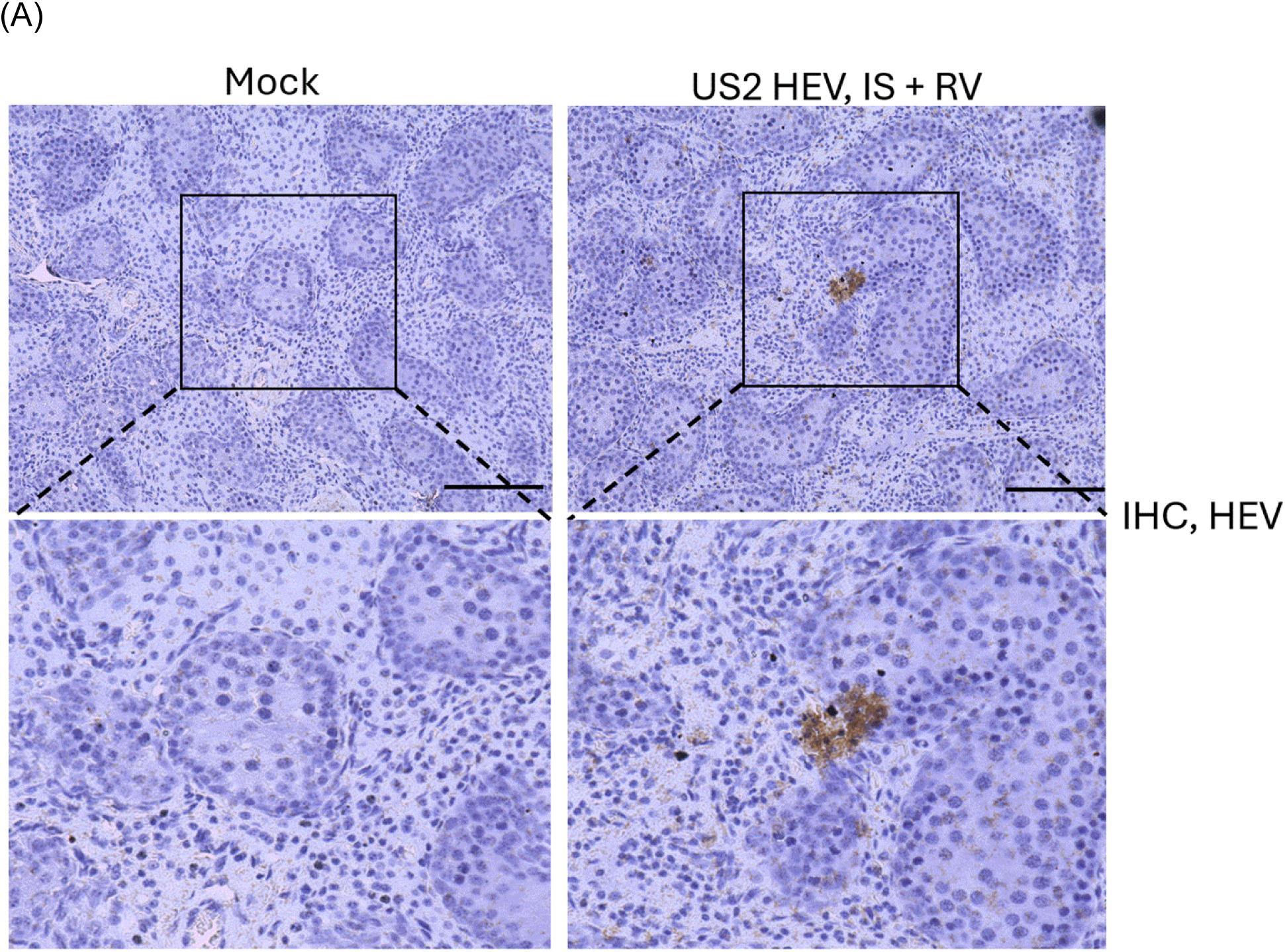
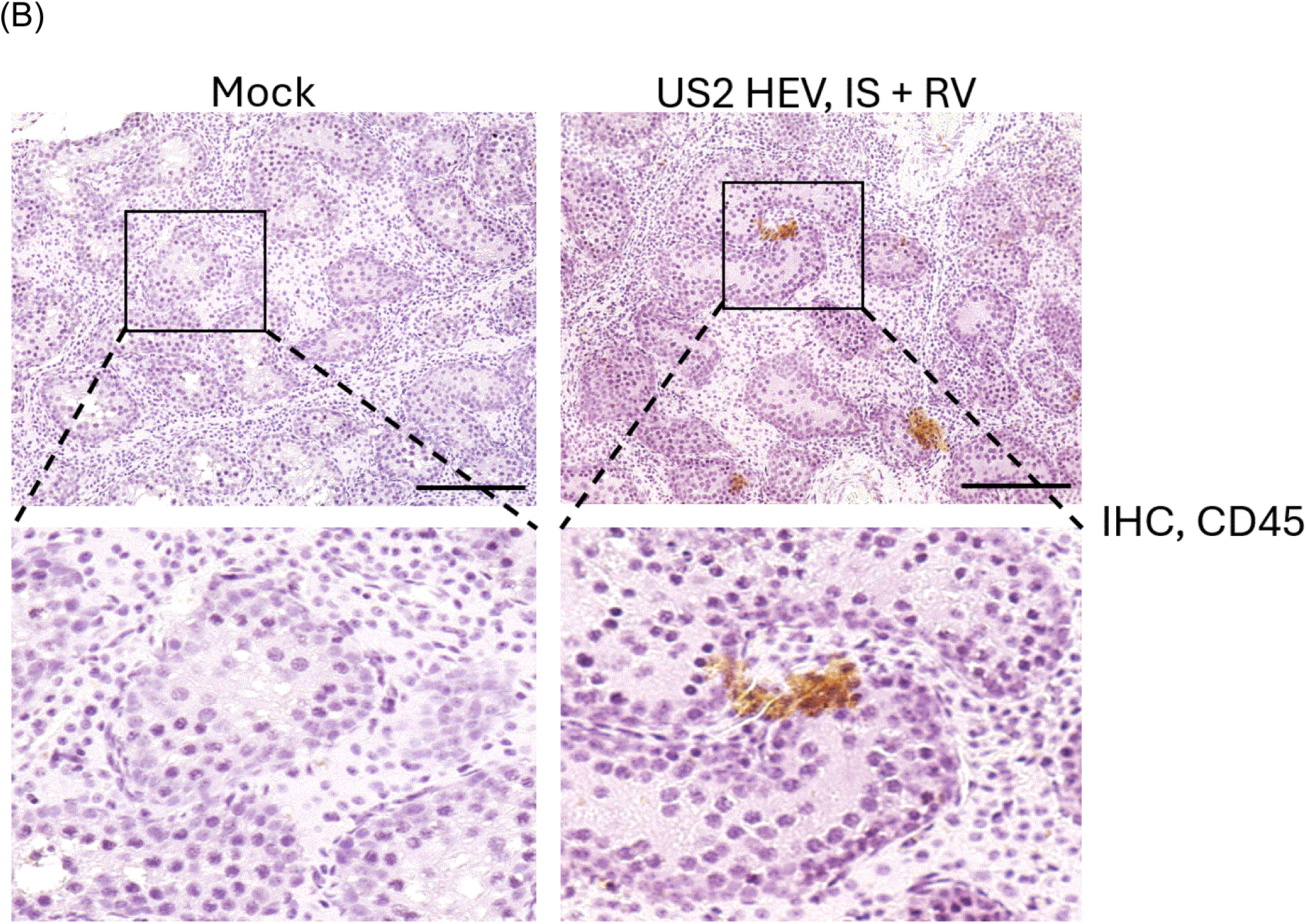
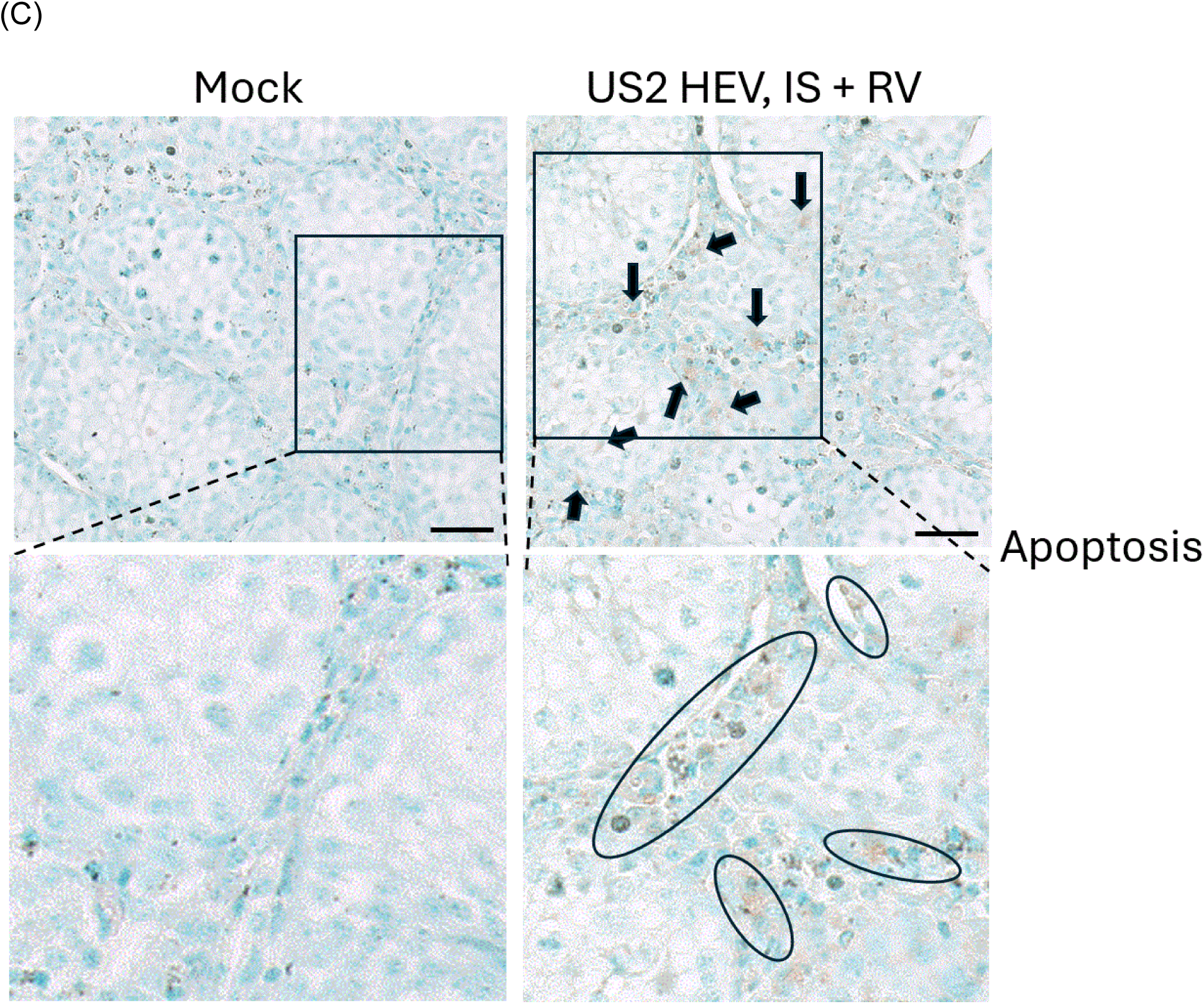
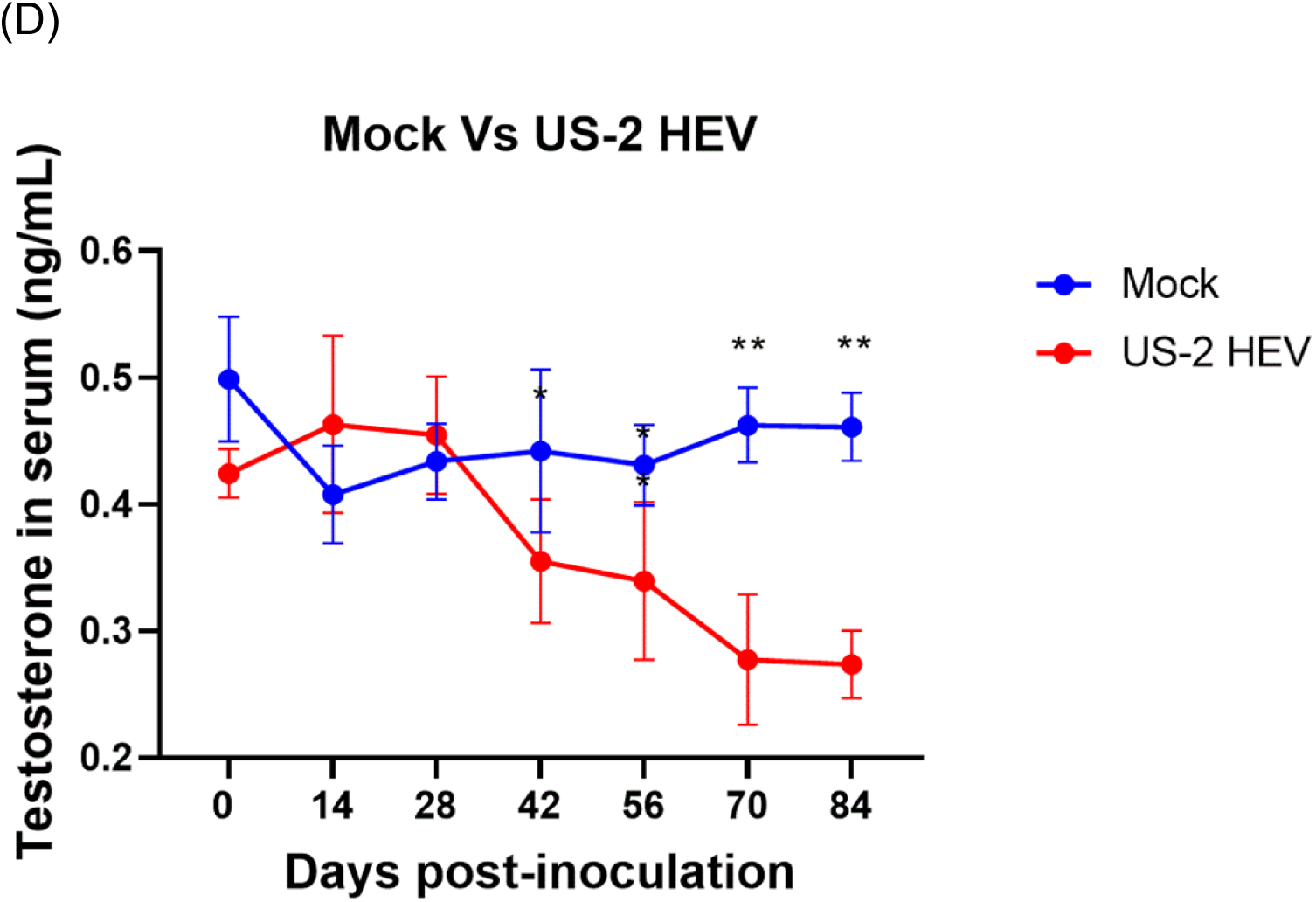
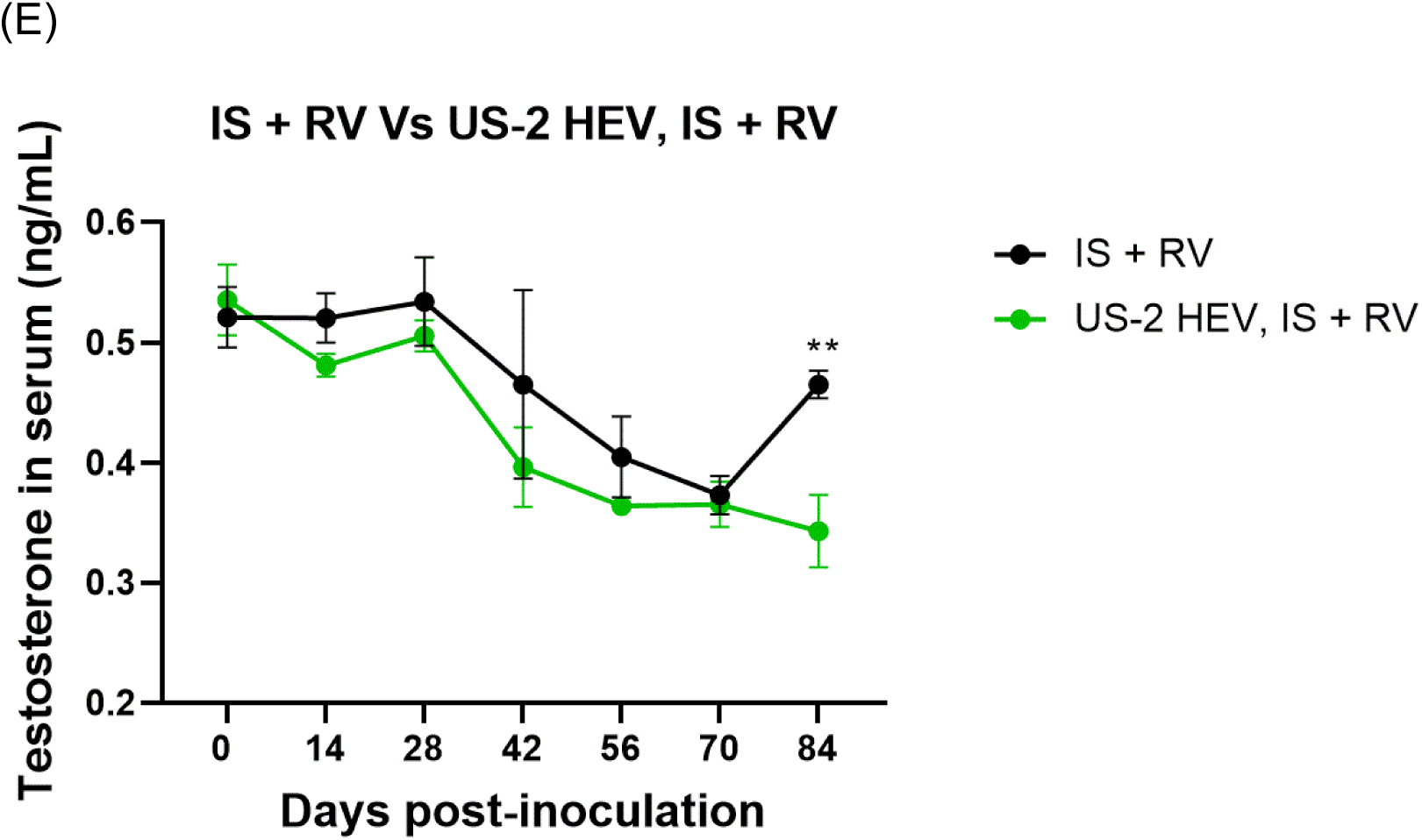
Hepatitis E virus replication, CD45+ leukocyte infiltration and apoptosis in blood testis barrier (BTB). (A) Immunohistochemical (IHC) staining of testis demonstrating the HEV open reading frame (ORF) 2 capsid protein and infiltration of CD45+ leukocytes (B) at the BTB in US-2 HEV infected, immunosuppressed (IS) and ribavirin (RBV) treated pigs. (C) TUNEL assay demonstrating apoptosis at the BTB of US-2 HEV infected, immunosuppressed (IS) and ribavirin (RBV) treated pigs. (D) Serum testosterone levels measurement between mock and US-2 HEV infected pigs demonstrates significant decrease in the virus infected group on day 70 and 84. (E) Serum testosterone level between immunosuppressed (IS) + ribavirin (RBV) and US-2 HEV infected, IS and RBV treated pigs were similar except on day 84.

### HEV replication in the male accessory glands

To evaluate the contribution of the male accessory glands in HEV reproductive pathogenesis, persistence, and contribution to semen, we utilized immunocompetent and immunocompromised pigs. HEV antigen was demonstrated in the prostate gland, Cowper’s gland, seminal vesicle, and epididymis on day 84 post inoculation (Fig. 5A, 5B). HEV viral RNA was higher in the prostate gland, followed by seminal vesicle, and Cowper’s gland in immunocompetent pigs compared to chronically infected immunosuppressed pigs (Fig. 5C). Moreover, we saw less HEV antigen in the immunocompromised pigs when compared to immunocompetent pigs. Higher HEV RNA was observed in the prostate gland in immunosuppressed pigs when compared to other accessory glands (Fig. 5C). Mock infected pigs and mock infected, immunosuppressed and ribavirin treated pigs did not demonstrate any HEV in the male accessory glands. Infiltration of CD45+ leukocytes was highly evident in the prostate gland, seminal vesicles, and epididymis of the US-2 HEV infected pigs on day 84 post inoculation. In contrast, US-2 HEV infected, immunosuppressed and ribavirin treated pigs demonstrate less infiltration of the CD45+ leukocytes on day 84 post inoculation (Fig. 5D, 5E). Prostate gland and seminal vesicle in the US-2 HEV infected, immunosuppressed and ribavirin treated pigs demonstrated less infiltration of CD45+ leukocytes. A TUNEL assay revealed higher apoptosis in the prostate gland, seminal vesicles, and epididymis of US-2 HEV infected pigs than in the US-2 HEV infected, immunosuppressed and ribavirin treated pigs on day 84 post inoculation (Fig. 5F, 5G).

**Figure 5.**
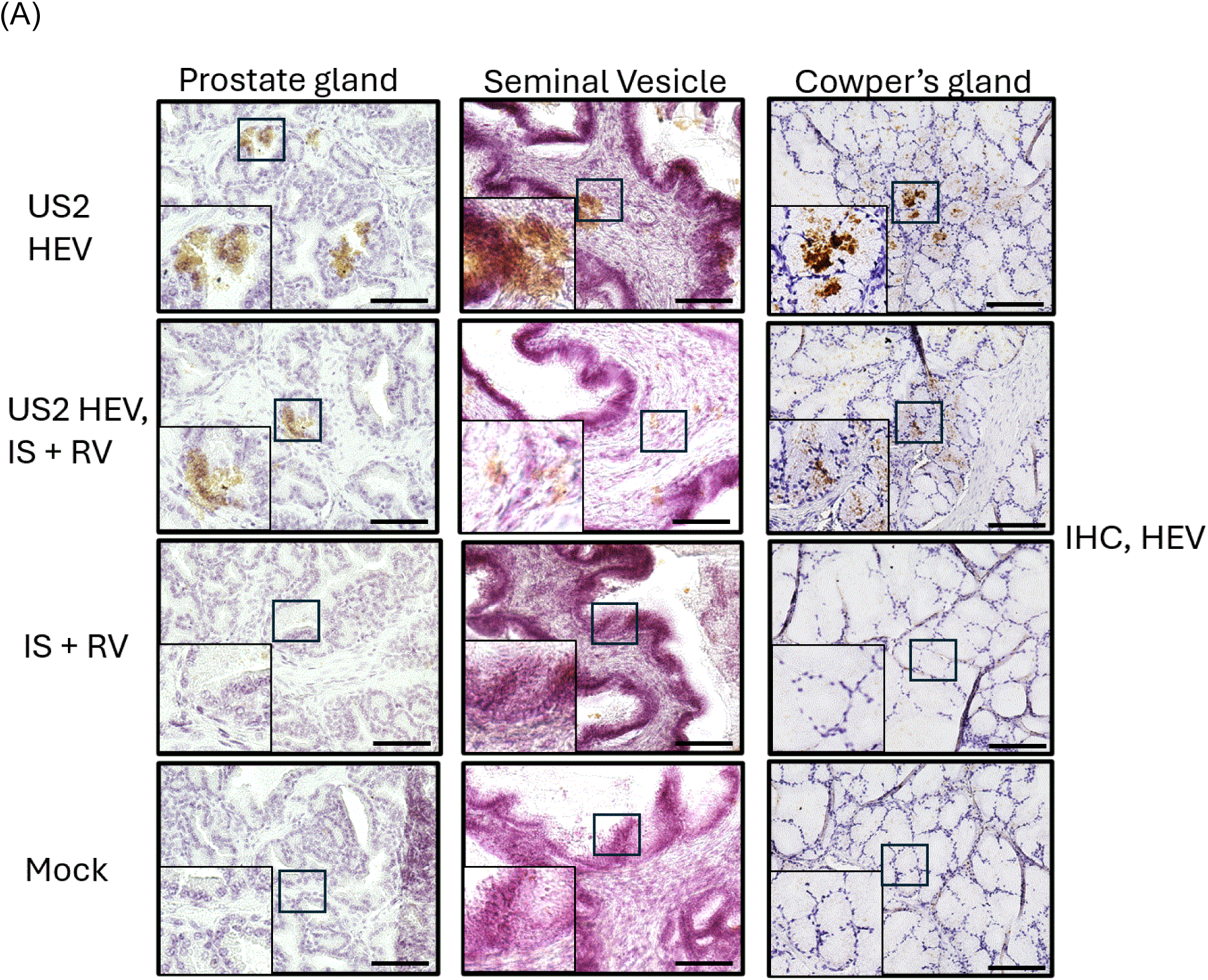
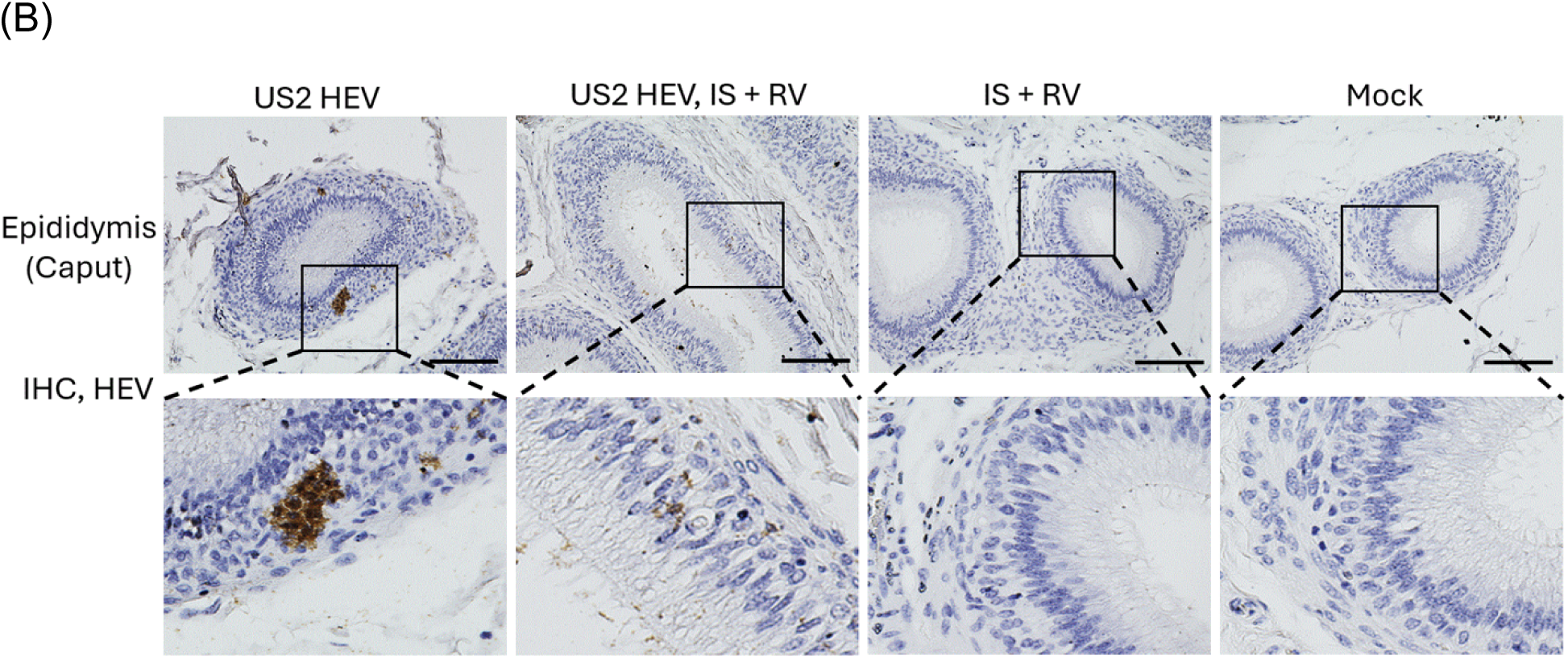
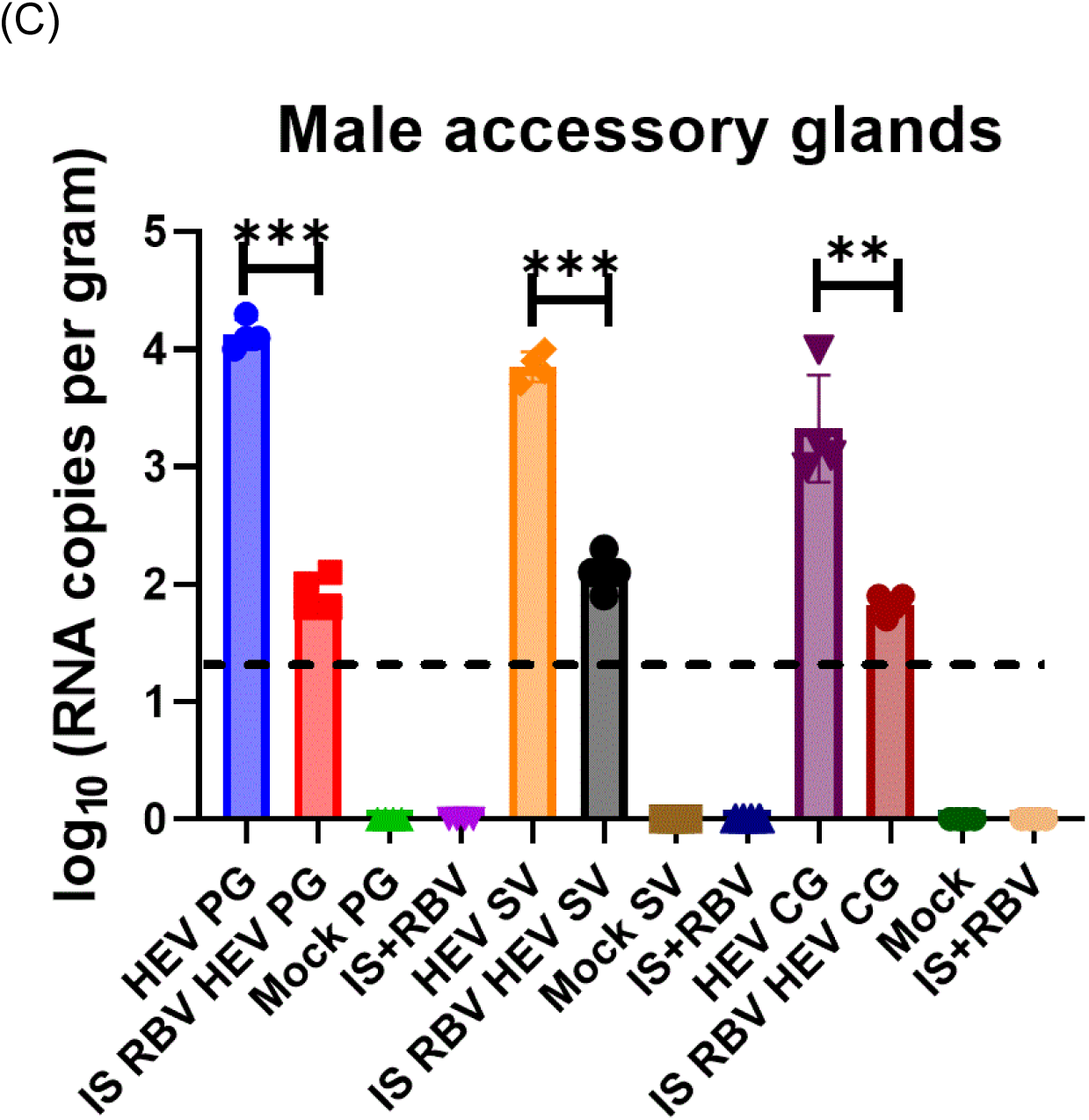
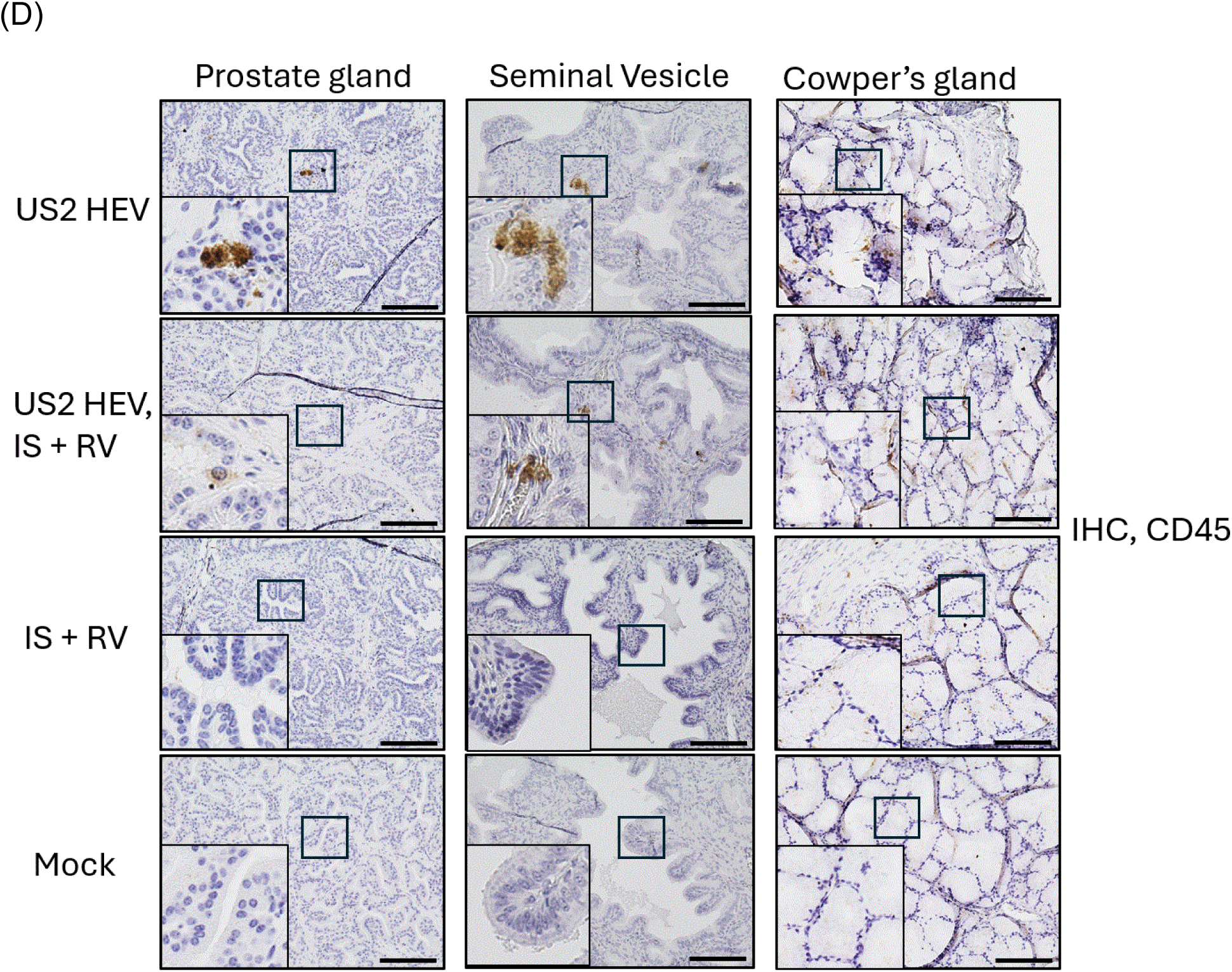
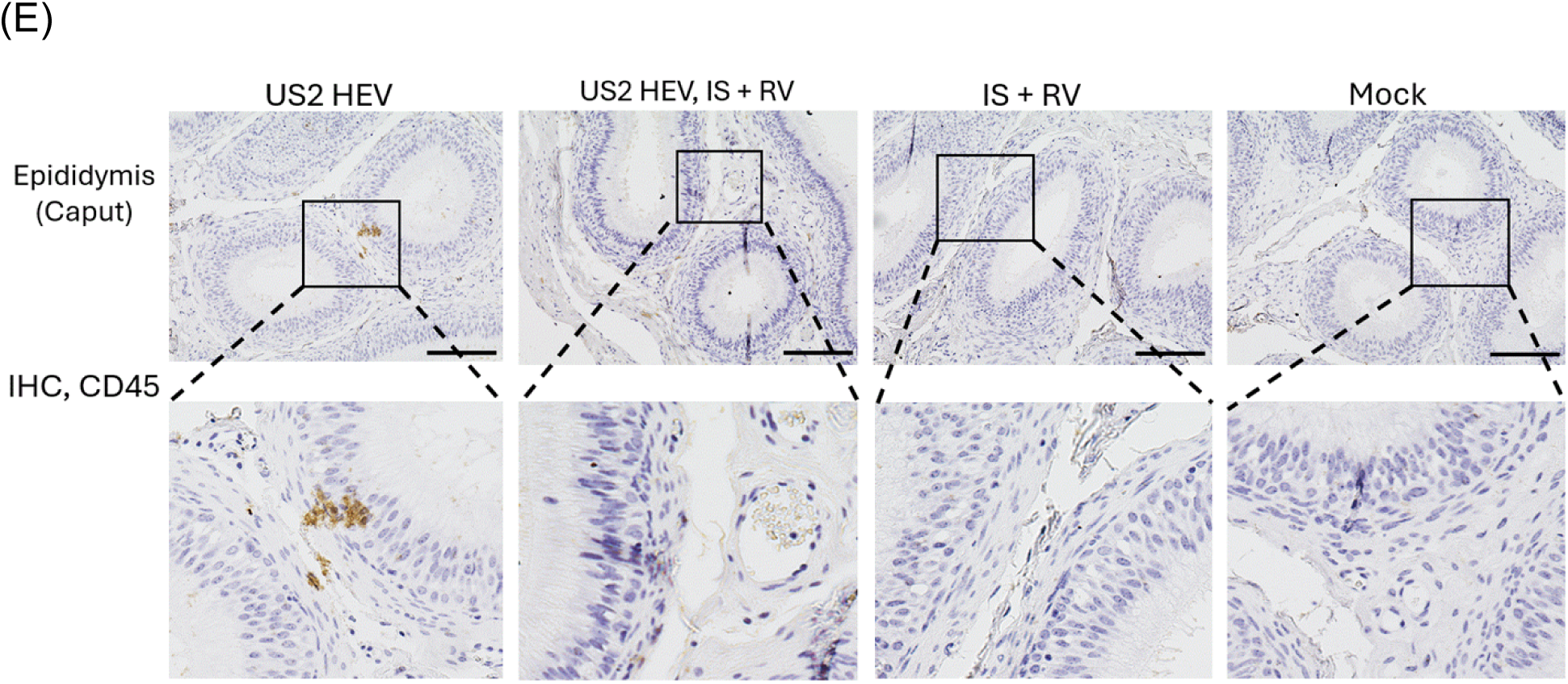
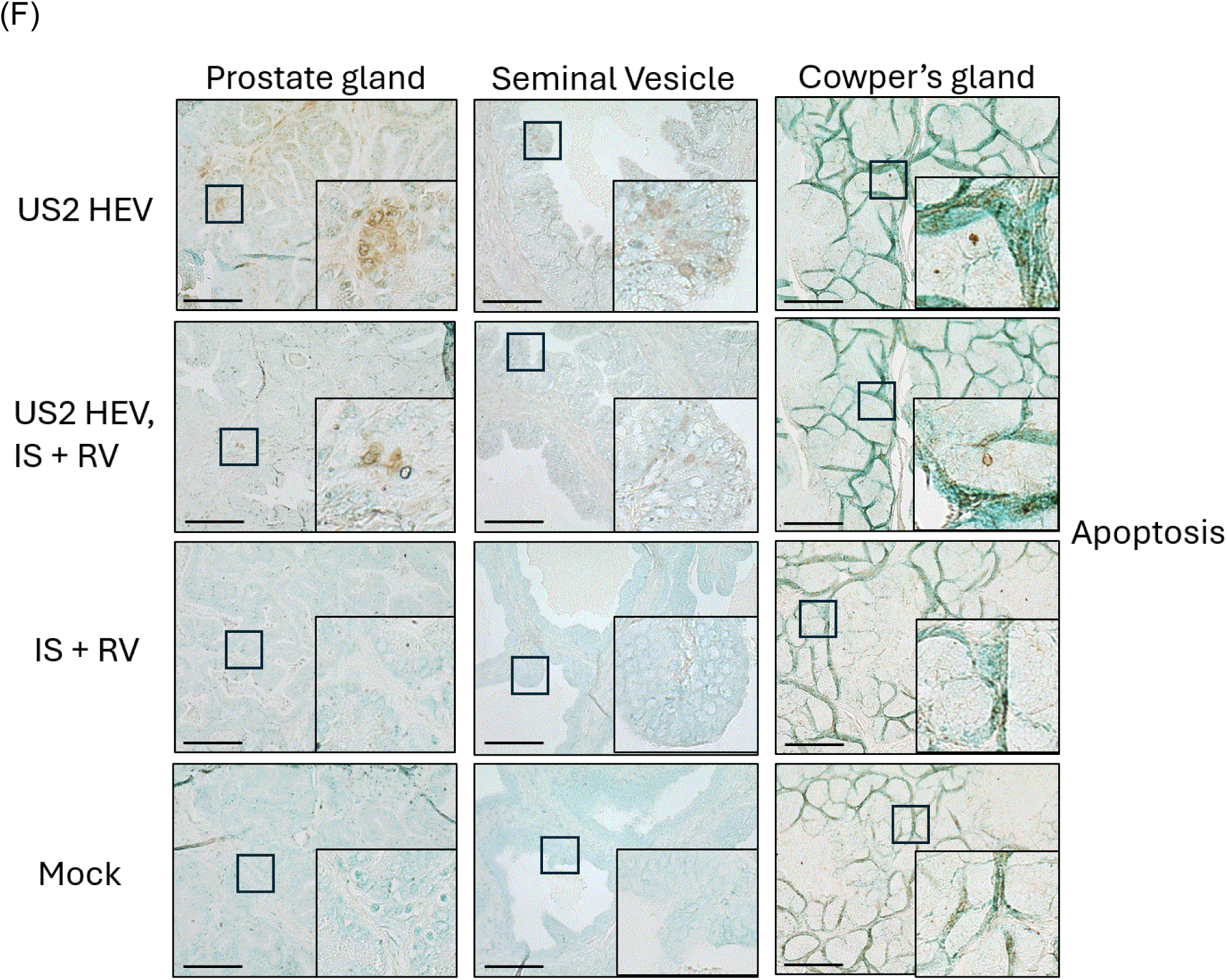
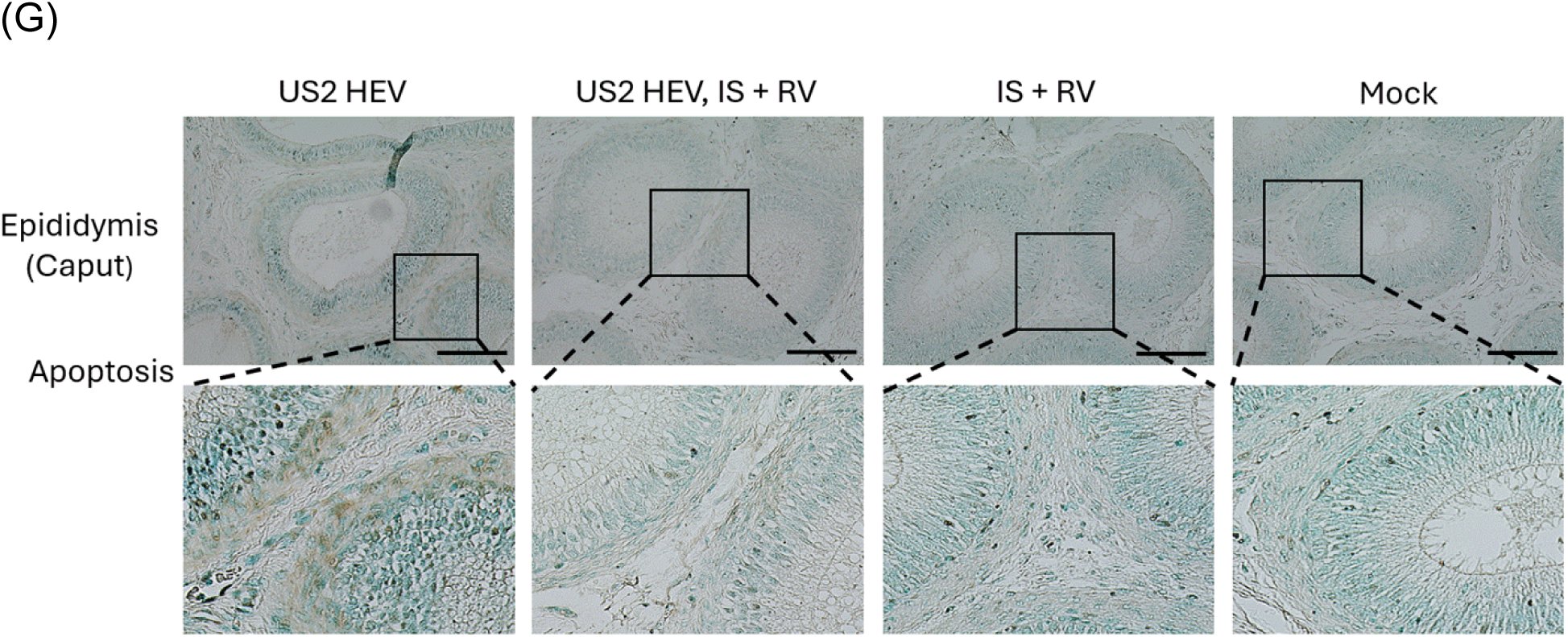
HEV infection and apoptosis in the male accessory glands. (A) Immunohistochemical (IHC) staining of prostate, seminal vesicles, Cowper’s gland and epididymis (B) demonstrating the HEV open reading frame (ORF) 2 capsid protein. (C) Viral RNA loads in the prostate gland, and seminal vesicle of mock, and immunosuppressed (IS) + ribavirin (RBV) and US-2 HEV infected, IS and RBV treated pigs. The dotted line represents the cut-off value demonstrating the background. ** indicates p < 0.01, *** indicates p < 0.001. (D) Infiltration of CD45+ leukocytes in prostate, seminal vesicle, Cowper’s gland and epididymis (E) between all four groups. (F) TUNEL assay demonstrating apoptosis in the prostate, seminal vesicle, Cowper’s gland and epididymis (G) between all four groups.

## Discussion

The presence of RNA virus in the head (26) and tail region (27) of sperm cells has been demonstrated for ZIKA virus revealing its ability and mode of sexual transmission in humans (28). The presence of infectious HEV in the sperm head is a very interesting finding highlighting a potential for sexual transmission of HEV (8). Our recent findings have shifted the paradigm of hepevirus-host interaction and highlight the need to understand possible sexual transmission of HEV.

Here, we demonstrate high viral RNA copies in sperm cells from immunosuppressed, ribavirin treated pigs. Our study necessitates an understanding of viral persistence in sperm cells after symptomatic and asymptomatic HEV infections. Infectious HEV seen in the sperm cells collected on day 84 suggests the need to understand the length of infectious virus persistence in sperm cells. Recent reports from chronically infected patients also demonstrated infectious HEV particles in the ejaculate (7) suggesting immunosuppressed pigs can be used as an efficient model to study the HEV effect on sperm health. Other viruses such as Zika and Ebola have been studied for sexual transmission between humans and have demonstrated a long-lasting excretion of viral RNA in the semen (29, 30). Zika virus has also been shown to cause a decrease in sperm motility and morphology in humans (31). A decrease in sperm motility has been reported in HEV infected animals (32, 33) and associated with infertility in humans (34). Similarly, our study demonstrated a decrease in the progressive motility of sperm cells in immunosuppressed chronically infected pigs. Previously, we reported 19% of the sperm cells associated with HEV ORF2 antigen in US-2 HEV infected pigs with high viral RNA titers. Interestingly, in the chronically infected, immunosuppressed and ribavirin treated pigs, only 7% of the sperm cells were positive for HEV ORF2 and showed comparatively lower viral titers. We speculate ribavirin treatment to be one of the major factors directly associated with our observed decrease of the virus in the sperm cells. Ribavirin is known to cross the BTB in humans (35) and rats (36) and thus we hypothesize that it may be playing a crucial role affecting the viral replication in the testis of pigs. Future studies need to be directed towards understanding the transport of ribavirin across the BTB in pigs and its effect in HEV quasispecies formation and overall impact in spermatogenesis.

Multiple viruses disrupt the BTB by various mechanisms. For instance, zika virus infection affects the permeability of the BTB through matrix metalloproteinase 9 (MMP9) mediated degradation of tight junction proteins and type IV collagens that are involved in the maintenance of BTB (37). HIV-1 disrupts the BTB permeability by utilizing its Tat protein transduction domain (38). Mumps and SARS-CoV-2 disrupt BTB via decreasing the expression of junctional proteins such as occludin, claudin-11 (39, 40). In our study, we demonstrate the damage in the BTB integrity during HEV infection. Apoptosis and infiltration of CD45+ leukocytes were evident at the BTB of chronically infected pigs. A recently published study suggested HEV possessed the ability to infect Sertoli cells and alter the cytokine milieu (41). Thus, we hypothesize that HEV disruption of the BTB leading to apoptosis could be either due to down expression or degradation of the junction proteins. It would be interesting to study the effects of the HEV viral protein in the expression of the BTB junctional proteins. Future research understanding the mechanism of BTB disruption by HEV is warranted.

Leydig cells are known as the primary source of androgens (42) and are located just adjacent to the BTB (43). Decreases in serum testosterone are directly related to a decrease in the numbers of Leydig cells (44). In our study, US-2 HEV infected pigs showed a decrease in serum testosterone levels on day 70 and 84 indicating a decrease in Leydig cells. Testosterone concentration has been demonstrated to affect spermatogenesis. In the absence of testosterone signaling: a) mature spermatozoa are retained within the seminiferous tubules and phagocytized by Sertoli cells (45, 46), b) BTB tight junction protein assembly rate is decreased (47), c) the amount of connexin between the Sertoli cells and male germ cells is reduced, leading to premature separation of round spermatids from Sertoli cells (48). In our HEV infected pig model, abnormal sperm morphology was documented in addition to apoptosis at the BTB. We speculate that HEV infection in the testicular tissues and significant decrease in the serum testosterone concentration causes disturbance in testicular homeostasis leading to decreased progressive motility as reported in our study.

In general, infection of the reproductive system activates the innate immune system. This response can disturb the testis-specific restricted immunological response leading to leukocyte infiltration and resulting in oxidative stress responsible for the damage to the testicular tissues and compromising the BTB integrity (49, 50). Infiltration of CD45+ cells were observed in the chronically infected immunosuppressed pigs. CD45+ lymphocyte infiltration is directly associated with abnormal sperm morphology and early apoptosis of the sperm cells (51, 52). Infiltration of CD45 in HEV infected mice testis has been attributed to the breakage of the BTB caused by HEV infection (15). Severe inflammation as indicated by CD45+ cell infiltration could be playing an important role in disruption of immune privilege leading to reduced sperm motility, and increased sperm abnormalities among HEV infected males. A long-term study demonstrating the viral clearance, status of BTB, immune cells and sperm quality report will be needed to understand if the resolution is partial or full after HEV infection.

Sexual transmission of HEV in humans has not yet been documented. We speculate this may be due to male partners of females demonstrating HEV infection are not routinely screened for HEV in their semen samples. Considering, the presence of viral RNA in chronic patients (5), viral RNA in the semen of infertile patients (53) and experimental studies demonstrating the infectious HEV in the ejaculate (7) and sperm head (8), further research is needed in larger cohorts to determine the contribution of a sexual route of transmission in human patients. Screening of male patients in HEV endemic areas is very much essential to determine the effect of HEV on sperm quality and reproductive health disorders.

Association of HEV with the female reproductive system has been studied for decades to understand the pathogenesis behind pregnancy mortality, but the mechanism by which HEV alters male fertility still awaits in-depth explanation. Recent literature suggests a new dynamic of HEV to the male genital system. HEV replication in human Sertoli cells (54), association with the sperm head in the pig model (8), and transmission to the liver through the intravaginal route in the rabbit model (55) necessitates study of the nature and mechanisms of HEV host interactions in the male genital tract. Semen is a mixture of sperm cells and secreting fluids, thus the presence of virus in the semen can be due to viral replication in the male accessory glands (56). Interestingly, the most studied tissues (testis and epididymis) of the male reproductive tract only contribute 10% to the semen volume (57). The major portion of the semen is contributed by the seminal vesicles (65-75%) and the prostate gland (25-30%) (57). Sperm parameters are directly influenced by the infection in the male accessory glands. Measuring the specific secretary products of these glands in the seminal fluid will allow us to understand the effect of one gland in comparison to others (13). However, they have not been studied well for their role in viral replication. The most studied RNA virus, HIV, has been demonstrated to infect seminal vesicles both in vitro and in vivo (58). Interestingly, infected seminal macrophages persisted longer than viremia demonstrating the seminal vesicle as a potential reservoir (58). Simian immunodeficiency virus (SIV) studies in macaques demonstrated higher infection in prostate and seminal vesicles followed by the epididymis and testis (59). Similarly, long term presence of Zika virus in the semen of vasectomized men (60) and reported sexual transmission of Zika from a vasectomized male to his partner (61) strongly suggests the importance of the prostate and seminal vesicles as a potential reservoir for sexual transmission. Such studies highlight the importance of accessory glands in the persistence of viral infection. In our study, HEV has been demonstrated in the male accessory glands, mainly prostate, seminal vesicle and epididymis of pigs harvested on day 84 post inoculation, suggesting HEV replication in these accessory glands. It would be interesting to study whether HEV can still be present in seminal fluids after vasectomy. These studies would further define the role of the accessory glands in the contribution of HEV in the semen. Additionally, the presence of leukocytes in the male accessory glands of pigs is a sign of infection and/or localized inflammatory response. It would be interesting to investigate the leukocytospermia and HEV infection associated effects in the sperm quality and overall reproductive health. A positive association between leukocytospermia, ROS, and sperm DNA fragmentation has been described (62). We speculate that the presence of leukocytes in the male accessory glands and testis could be a contributing factor to low progressive motility seen in HEV infected pigs.

The virus shedding in the semen also depends on viral attachment to the specific receptors present in the spermatozoa. In the case of HIV, alternative receptors such as heparan sulfate and Ga1AAg (glycolipid-related glyceramide) have been demonstrated to play an important role in the capture of HIV (63). Interestingly, one of the receptors defined for HEV is heparan sulfate (64). Studies understanding if heparan sulfate of sperm cells have a role in the capture of HEV will be an interesting topic. As HIV has been studied for decades, semen has been considered as the main vector for the HIV-1 dissemination. Three major sources of infectious HIV have been reported: free virions, infected leukocytes, and spermatozoa-associated virions (65). Future studies characterizing the presence of HEV in the semen following the similar differentiation is essential to understand the duration of infectious HEV present in the semen.

## Materials and Methods

### Animals, immunosuppressive drugs, and virus

Eight pigs (US-2 HEV infected, n1 = 4, mock infected and drugs treated, n2=4) were administered immunosuppressive (IS) drugs (Cyclosporine: 10 mg/kg/day, Prednisolone: 4 mg/kg/day and Azathioprine: 2 mg/kg/day) following the established regimen from the Meng laboratory (23). These eight pigs were also administered ribavirin (RBV): 80 mg/day. At six-weeks-old, male conventional pigs in the US-2 HEV infected group were anesthetized and infected via ear vein inoculation with (2 × 10^8^) viral RNA copies (log_10_/ml) of gastrointestinally-derived genotype 3 US-2 (human) hepatitis E virus (66). The remaining four pigs only continued with the immunosuppressive drugs and ribavirin throughout the study. Four pigs (n3 = 4) were used as a negative control in the study and the other four (n4 = 4) were only infected with US-2 HEV. This route of infection mimics what might occur in the case of human transfusion acquired HEV infection.

### Collection of semen from US-2 HEV inoculated pigs

Testis were collected from pigs at 84 days post inoculation (dpi) (age of pigs = 126 days old). The testis was extracted during necropsy by a veterinarian and stored in a cooled box. Semen from the epididymis was harvested by aspiration and flushing (67). The procedure was repeated for both testis from the same animal and the extracted semen was pooled.

### Examination of semen parameters

Semen samples were examined within 15 minutes after collection following the 2010 World Health Organization laboratory manual for the examination of human semen (68). Motility and morphology of sperm cells between the groups were studied. In brief, 200 spermatozoa per replicate were examined at 200x magnification. Motility in a sperm cell was studied using three categories: progressive (PR), non-progressive (NP), and immobility (IM). PR refers to a percentage of a sperm’s progressive motility (moving actively, either linearly or in a circle, regardless of speed). NP refers to non-progressive motility (all other patterns of motility with absent progression). IM refers to complete immobility.

### Sperm cell separation from the semen

Semen is a mixture of seminal plasma fluid and sperm cells (69). Sperm cells were separated using phosphate buffered saline (PBS) in a ratio of 1:10 (volume/volume), and gently mixed to remove the sperm cell clumps. Supernatant was collected and stored at −80°C post centrifugation at 1500 rpm for 10 min. To remove any remnants of seminal plasma, the sperm cell pellet was washed once with 900 µL of PBS, centrifuged, and the supernatant was discarded. The remaining pellet was carefully resuspended with another 900 µL of PBS (27).

### Immunofluorescence staining of sperm cells

To identify hepatitis E virus antigen in sperm cells, 15 µL of the sperm suspension was added in duplicate to Superfrost Plus Slides (Fisher) dried and fixed with paraformaldehyde solution (4% in PBS) at 4°C for 5 min, followed by two rounds of washing in PBS for 5 min at room temperature. For the detection of HEV antigens, 1:200 rabbit anti-HEV open reading frame 2 (ORF2, encodes structural capsid protein of HEV) polyclonal antibody (Cocalico Biologicals from in-house prepared ORF2 protein lacking the first 111 amino acids produced in E coli, denatured, and polyacrylamide gel purified) was added to each slide and incubated for 45 minutes at room temperature followed by three washings in PBS for 5 min each. Translation of ORF2 occurs from subgenomic mRNA, is synthesized during the later stages of replication, and thus an indicator of active viral replication (70, 71). Goat anti-rabbit IgG H&L (Alexa Fluor 594; Abcam) (1:500) was used as the secondary antibody and the slides were incubated for 30 minutes at room temperature. 30 µL DAPI (4’,6-diamidino-2-phenylindole) was added as a nuclear counterstain for fluorescence microscopy. Slides were imaged using a Keyence microscope at 40x magnification. A sample was considered positive if at least one cell with clear sperm like morphology contained red fluorescence. All specimens were examined in duplicate.

### Flow cytometry analyses

400 µL of the sperm cell suspension was used for flow cytometry quantification of HEV positive cells. Fixed cells were probed with primary antibody – rabbit anti ORF2 HEV diluted 1:200 in PBS for 30 min at 37°C. Cells were then washed twice with PBS and incubated with secondary antibody – goat anti-rabbit phycoerythrin (Life Technologies) diluted 1:500 in PBS for 30 min at 37°C. Cells were then washed and resuspended in 200 µL of PBS. Fluorescence was analyzed for 100,000 events using a flow cytometer (BD Accuri C6 plus). Histogram plots were compared between the infected and mock infected sperm cells.

### Infectivity assay

450 µL of the sperm cell suspension (∼4 × 10^6^ viral RNA copies) was used to assess the infectiousness of HEV derived from sperm cells. Human liver cells (Huh7 S10-3) were used for the study (72). Three repeated freeze and thaw cycles were done with the sperm cell suspension. Cell lysates including both the cell debris and virus were subjected to high-speed centrifugation (10,000 x g for 5 min) which separates the cell debris. The collected supernatant was used to inoculate Huh7 S10-3 liver cells. Eight hours post inoculation, the cell culture media was removed, and fresh media was added. Forty-eight hours post inoculation, cells were passed 1:3 and 72 hours later were fixed for immunofluorescence assays.

### Indirect immunofluorescence

At 5 dpi, human liver cells were fixed and permeabilized with PBS plus 0.5% tween 20 (PBST). 5% non-fat dried milk (Sigma-Aldrich) in PBST was used to block non-specific antibody binding. Immunostaining of HEV ORF2 capsid protein was performed using 1:200 rabbit anti-truncated ORF2 HEV. A fluorescently labelled goat anti-rabbit IgG H&L secondary antibody (Alexa Fluor 594; Abcam), 1:500, was used. DAPI was used to stain the nucleus. Images were optimized for intensity enhancement using Coreldraw 2019 applied to all images equally.

### RNA extraction and RT-qPCR

RNA was extracted using TRIzol reagent (Invitrogen). Extracted RNA from serum, rectal swabs, sperm cells, and seminal fluid was quantitated using reverse transcriptase quantitative polymerase chain reaction (RT-qPCR). Primers US-2 HEV F, 5′-GGTGGTTTCTGGGGTGAC-3′and US-2 HEV R, 5′-AGGGGTTGGTTGGATGAA-3′, and probe 5’-FAM-TGATTCTCAGCCCTTCGC-Dabcyl-3’ were used for the detection of US-2 HEV. A 10-fold serial dilution of US-2 HEV RNA (10^7^ to 10^1^ copies) was used as a standard for quantification of the viral genome copy numbers.

### Immunohistochemistry (IHC)

Formalin-fixed tissue sections were tested by IHC for the detection of HEV, as previously described with slight modifications (73). The rabbit anti-HEV-ORF2 antibody was used as the primary antibody and a horseradish peroxidase conjugated anti-rabbit antibody (BioGeneX) was used for visualization as brown staining. Stained tissues were counterstained with hematoxylin. In addition, the tissues were also tested by IHC for the detection of CD45 expressing cells. Mouse anti pig CD45 (Bio-RAD) was used as the primary antibody and a horseradish peroxidase conjugated anti-mouse antibody (BioGeneX) was used for visualization as brown staining.

### Testosterone assay

Serum samples from pigs were tested for the testosterone concentration on day 0, 14, 28, 42, 56, 70 and 84 post inoculation using a pig testosterone enzyme-linked immunosorbent assay (ELISA) kit (CUSABIO, CSB-E06796p). Briefly, 50 uL of the sample was added in a well followed by 50 uL of HRP-conjugate and 50 uL of antibody. The samples were incubated for 1 hour at 37C after mixing. Later the solution was aspirated, discarded and wells were washed three times with the wash buffer. Then, 50 uL of substrate A and substrate B was added to each well, mixed and the plate was incubated for 15 minutes at 37C in the dark. The stop solution was added, and the absorbance was read at 450 nm using a microplate reader (FiltermaxF5, Molecular Devices).

### Terminal deoxynucleotidyl transferase dUTP nick-end labeling (TUNEL) assay

Apoptotic cells in the testicles of pigs were detected by using One Step Apoptosis Assay Kit (Abcam, ab206386) according to manufacturer’s directions. Briefly, tissue sections were deparaffinized, hydrated, blocked with proteinase K for 15-30 min, and washed with PBS. After quenching with the 3% hydrogen peroxide, tissue specimens were covered with TdT equilibration buffer and incubated for 30 min followed by labelling reaction using the TdT labelling reaction mix. The tissue specimens were treated with DAB solution followed by Methyl Green counterstain solution.

### Statistical analysis

Analyses of independent data were performed by Student’s unpaired two-tailed t test and two-way ANOVA using GraphPad Prism 9.4.1. p < 0.05 was considered statistically significant.

## Acknowledgements

We thank the animal care staff (Megan Strother, Sara Tallmadge) for their contribution to animal care and handling training.

## Author contributions

K.K.Y. and S.P.K. designed the study. J.H. contributed to collecting testicle samples from the pigs. K.K.Y., T.L., M.B., and S.P.K. contributed to in vitro experimental designs and data analysis. K.K.Y. and S.P.K. wrote the manuscript and prepared the figures. K.K.Y., T.L., P.B., S.K., C.L., J.H., and S.P.K., contributed to in vivo experimental design. All authors reviewed the final manuscript and approved the submission. All data are presented in the paper.

## Competing interests

We declare no competing interests.

## Notes

### Competing Interest Statement

The authors have declared no competing interest.

